# Zero-inflated Joint Species Distribution Models for improved partial-correlation network inference from community data

**DOI:** 10.1101/2025.07.24.666553

**Authors:** Jeanne Tous, Julien Chiquet, Amy E. Deacon, Ada Fontrodona-Eslava, Douglas F. Fraser, Anne E. Magurran

## Abstract

1. A long-term goal of community ecology has been to decipher the mechanisms that shape the spatio-temporal organization of species communities. Understanding these processes is critical to predicting the responses of ecological communities to environmental change. To this end, Joint Species Distribution Models (JSDMs) offer statistical tools to analyze community data, identify the impact of abiotic factors on them and study inter-species correlations in their distributions. In particular, the JSDM-inferred partial-correlation networks allow one to identify direct links between species that can help decipher the mechanisms that shape their joint distribution.
2. Community data based on species counts often contains numerous zeros. However, not accounting for these zeros in a data set is known to hinder parameter inference. We investigate this issue in the context of JSDMs, and ask what impact it can have on the inference of partial-correlation networks.
3. We propose a novel JSDM, the ZIPLN-network model, based on the PLN-network (Poisson log-normal network) and ZIPLN (Zero-Inflated Poisson log-normal) model, which models count data while including zero-inflation and infers a partial-correlation network. Using simulated data, we compare the results obtained by this model with existing JSDMs in terms of association network inference from abundance data containing structural zeros. We then illustrate the ZIPLN-network approach using real data from tropical freshwater fish communities.
4. Simulations show that zero-inflation can significantly bias the inference of partial-correlation networks from community data and that the ZIPLN-network model efficiently counterbalances these effects. The ZIPLN-network approach is widely applicable to community data, delivers ecologically-insightful analyses, helps distinguish amongst potential mechanisms, and aids better understanding of community assembly rules. We provide guidance for getting started with our approach.

## 1. Introduction

Species community data typically describe the distribution of a set of species, in a collection of sites, and can be based on presence-absence, biomass, environmental DNA or counts of individuals at each location [Ovaskainen and Abrego, 2020]. A long-term goal of community ecology has been to infer the processes that shape the spatio-temporal structures of species communities from such observational data [Forbes, 1907, Connor and Simberloff, 1979, Diamond, 1975, Gotelli and Graves 1996, Ovaskainen et al. 2010]. Perspectives developed to study this question [Vellend, 2010], include descriptive, pattern-based approaches [Whittaker, 1962] and the niche theory [Grinnel, 1917; Hutchinson 1957; Soberón, 2007]. A further approach is to study the three categories of mechanism shaping species distributions across different sites [D’Amen et al., 2017; Guisans and Thuiller, 2005; Poggiato et al., 2021] - abiotic factors through environmental filtering (whether the environment is suitable for a given species [Cadotte and Tucker, 2017, Kraft et al., 2015]), biotic interactions (how species’ presences influence one another) and dispersal (whether a species has been able to reach a site). Understanding how these mechanisms structure biodiversity patterns is critical for predicting ecosystem responses to environmental stressors, and for designing effective conservation strategies [Heinen et al. 2020]. However, the complicated ways in which these mechanisms operate, their intermingled influences, and the way their relative effects can depend on scale [Götzenberger et al. 2012, Viana and Chase 2019] make community data both information rich and difficult to interpret.

Multiple approaches have been developed to explore these questions. Early on, ordination methods [Bray and Curtis, 1957] were developed to represent multivariate community data over a reduced number of axes so as to identify main axes of variation in species community composition and relate them to environmental and geographical conditions; these were typically based on statistical methods such as Principal Component Analysis or Principal Correspondence Analysis [Gower, 1987, Legendre and Legendre, 2012]. Variation partitioning is a related approach that aims to quantify what fraction of the variation in community composition is attributable to environmental variables and to spatial auto-correlation [Peres-Neto et al., 2006, Smith and Lundholm, 2010].

Joint Species Distribution Models (JSDMs) [Pollock et al., 2014, Niku et al., 2019, Ovaskainen and Abrego, 2020, Tikhonov et al., 2020, Chiquet et al., 2021, Pichler and Hartig, 2021] are a more recent approach to this question.

JSDMs build on Species Distribution Models (SDMs). SDMs represent the relationship between the presence-absence or abundance of each species, and environmental covariates (such as temperature, precipitation, soil nature…) with, occasionally, the additional inclusion of biotic interactions [Poggiato et al., 2025, Staniczenko et al., 2017] or dispersal effects [Shipley et al., 2022].

Extending SDMs by modeling the distribution of a whole species community at once, instead of one species at a time, gives JSDMs a community ecology perspective, and supports the study of species co-occurrence patterns. This involves modeling of inter-species residual correlations after accounting for environmental and / or spatial predictors.

In their focus on inter-species correlations, JSDMs are part of a century-old approach in community ecology in which species co-occurrences are used to understand the underlying ecological processes [Forbes, 1907, Diamond, 1975, Blanchet et al., 2020]. Although it was long hoped that inter-species correlations would serve as proxies for biotic interactions, no such equivalence holds [Blanchet et al., 2020; Poggiato et al., 2021]; nonetheless they remain useful for generating hypotheses about underlying mechanisms (with biotic interactions being one possible explanation, but not the only one) [Gotelli et al., 2010; Ovaskainen and Abrego, 2020] and for improving predictions [Poggiato et al., 2021, Poggiato et al., 2025]

From a technical point of view, JSDMs generally model the response variable (the presence / absence or abundance of all the species on each site) as a multivariate random variable. Given a latent Gaussian variable, the components of this multivariate random variable are independent and each follow an appropriate distribution (e.g. Bernoulli for presence-absence, Poisson log-normal for species counts, log-normal for biomass). The parameters of this distribution usually depend on environmental predictors. The latent Gaussian variable allows for the modeling of a dependence structure, with its covariance matrix representing inter-species correlations [Pollock et al., 2014, Niku et al., 2019, Ovaskainen and Abrego, 2020, Tikhonov et al., 2020, Chiquet et al., 2021, Pichler and Hartig, 2021]. To reduce the number of parameters to be inferred in this matrix, different strategies exist. These include replacing the matrix with a lower-rank approximation using latent factors [Niku et al., 2019, Ovaskainen and Abrego, 2020], or using a penalized approach [Chiquet et al., 2021; Pichler and Hartig, 2021].

Two types of inter-species residual correlations can be considered in the output of a JSDM. The variance-covariance matrix of the latent Gaussian variable contains marginal correlations whereas its inverse, the precision matrix, contains partial correlations. The two matrices are related and one can, in theory, be obtained from the other, but when a JSDM resorts to a low-rank approximation of the variance-covariance matrix [Pollock et al., 2014, Niku et al., 2019, Ovaskainen and Abrego, 2020, Tikhonov et al., 2020, Pichler and Hartig, 2021], inverting it requires using statistical tricks, like regularization, and may yield a biased estimate of the precision matrix. Some JSDMs “natively” focus on precision matrices and partial correlations to model species association networks. They directly target the precision during the estimation procedure, using for instance a sparse prior to select strong direct associations encoded by the precision matrix [Chiquet et al., 2021]. This approach belongs to the framework of graphical models in statistics [Lauritzen, 1996, Whittaker, 2009] and is more widespread in microbial ecology [Kurtz et al., 2015; Schwager, 2017] than in community ecology.

In terms of interpretation, in JSDMs, residual marginal correlations (i.e., the entries of the variance matrix) indicate whether species co-occur more or less than predicted by the environmental covariates and other modelled covariates. Conversely, residual partial correlations (i.e., the entries of the precision matrix) represent conditional dependencies: the association that remains between two species once all the other species in the data have been accounted for. If an apparent association between species A and B arises solely through a third species C, the corresponding entry of the variance-covariance matrix is non-zero whereas that of the precision matrix is zero. Partial correlations are thus said to capture direct associations, in a statistical sense only, since they may still arise from unobserved species or unmeasured environmental covariates. Their study has been put forward as a promising tool in ecology, precisely because it separates associations mediated by other species from those that do not call for such an explanation [Harris, 2016; Popovic et al., 2019].

Our paper focuses on the inference of partial-correlation networks by JSDMs based on count data. While presence-absence data is more widely used, in particular because it is easier to collect, abundance (count-based) data give access to more detailed information and can help improve the understanding of the processes shaping ecological communities [Estrada and Arroyo, 2012, Sander et al., 2017], provided the collected data are reliable.

Count-based community data often contain numerous zeros [Heilbron 1994, Martin et al., 2005, Blasco-Moreno et al., 2019]. When the proportion of zeros exceeds what is predicted by standard distributions (Poisson log-normal, Negative-Binomial), the data set is said to be zero-inflated (with respect to a given distribution). Zeros in such data can be either false, random or structural [Martin et al., 2005, Blasco-Moreno et al., 2019]. False zeros result from observer’s errors (e.g. a poorly equipped observer missing an observation) or errors in the experimental design (sampling at the wrong place / time), random zeros arise from sampling variability, and structural zeros can be a consequence of environmental conditions, such as dispersal barriers.

It is widely appreciated that not accounting for zero-inflation in a data set can hinder parameter inference [Barry and Welsh, 2002, Martin et al., 2005], although, to our knowledge, there is no specific study of the way zero-inflation impacts species association networks (and specifically, partial-correlation networks) inference. Overall, when considering the presence of zeros in community data, while false zeros should be identified and removed when possible, specific methods are required to deal with other types of zeros [Blasco-Moreno et al., 2019]. Indeed, accounting for such zeros is a long-standing issue in the modeling of species abundances [Cunningham and Lindemayer, 2005; Nolan et al., 2022, Wenger and Freeman, 2008].

One strategy for dealing with this issue is the use of zero-altered models, also known as hurdle models [Mullahy, 1986]. These separately model zero and non-zero counts by first fitting a presence-absence model and then fitting an abundance model conditionally on presence. This is the approach used by the Hmsc model (Hierarchical modeling for Species Communities) to model extra zeros [Ovaskainen and Abrego, 2020]. A second strategy consists in directly modeling zero-inflation in an abundance model. Zero-inflated models consist of a mixture of two distributions, a Dirac zero distribution and a reference distribution to model abundance. In such a framework, a zero abundance may arise from either the zero distribution or the abundance distribution. This is the strategy used in the gllvm model (Generalized Linear Latent Variable Models) [Niku et al., 2019, Korhonen et al., 2025] and in the glmmTMB model [Brooks et al., 2017]. While both approaches to modelling excess zeros are closely related [Ovaskainen and Abrego, 2020], zero-inflated models allow an integrated analysis and appear more relevant when zeros also arise from the abundance distribution [Blasco-Moreno et al., 2019] - as is the case with a Poisson distribution.

Here we introduce the ZIPLN-network (Zero-Inflated Poisson log-normal network) model as a JSDM that models zero-inflated count data and uses a penalized method to infer a sparse network of inter-species partial correlations. It is based on the ZIPLN model [Batardière et al., 2024], a zero-inflated Poisson log-normal model with a latent variable but no specific focus on association network inference, and the PLN-network model [Chiquet et al., 2019], a Poisson log-normal model that focuses on association network inference by adding a l1-penalty on the latent variable’s precision matrix, in a LASSO-like approach [Friedman et al., 2008, Hastie et al., 2008] but that does not include zero-inflation. The novelty of the ZIPLN-network model consists in mixing these two approaches to build a zero-inflated PLN model with a focus on the inference of a sparse latent network of partial correlations.

In this study, we investigate the effects of different types of zero-inflation on the inference of partial-correlation networks by JSDMs. We examine to what extent zero-inflation can be detrimental to their retrieval, and study how different JSDMs can compensate for the effects of zero-inflation as well as how the ZIPLN-network model can help produce more informative analyses of zero-inflated community data thanks to its modeling of zero-inflation.

We run a simulation study to test to what extent zero-inflation impacts the quality of the inference of partial-correlation networks from virtual community data, and how accounting for zero-inflation can help mitigate this effect. To this end, we compare the results obtained with the ZIPLN-network, Hmsc and gllvm approaches.

We further exemplify the use of the ZIPLN-network model with an application to a species community dataset of tropical freshwater fishes sampled in the Caribbean island of Trinidad. This dataset contains numerous zeros, including structural ones owing to natural barriers such as waterfalls. We show that modeling these zero-inflation patterns filters out numerous statistical associations, and discuss how the resulting network sheds light on how the underpinning ecology can be interpreted.

Below, we first introduce the ZIPLN-network model and the methods we compare it to, before describing the simulation protocol. Next we provide an overview of the dataset used to illustrate the ZIPLN-network model. We finally present and discuss the results of both the simulation and real data studies, and mention possible improvements to the ZIPLN-network model.

## 2. Materials and methods

### 2.1 The effects of zero-inflation on correlation structures

While it is known that zero-inflation tends to impede the inference of a model’s parameters [Barry and Welsh, 2002, Martin et al., 2005], we do not know of any specific study in the context of JSDM-inferred association networks.

Statistically, zero-inflation is defined as an excess of zeros in a dataset compared to what a reference distribution could produce. A set of zero-inflated count observations is typically modelled as:

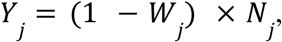

with *Y_j_* the number of individual counts of species j, *N_j_* a random variable following a given distribution (e.g., Negative Binomial or e-Poisson log-normal) and possibly related to environmental covariates, and *W_j_* following a Bernoulli distribution.

The focus of this study is the situation in which one wants to infer the correlation structure of variable *N* but only has access to observations *Y*. Taking the covariance term between the observations *Y_j_* and *Y_k_* for two species as:

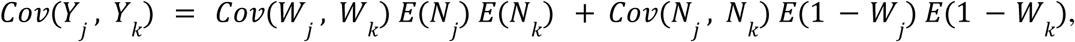

reveals that the zero-inflated variable *W* induces a false dependency expressed by the term *Cov*(*W_j_*, *W_k_*) *E*(*N_j_*) *E*(*N_k_*) and biases the true dependence by a factor *E*(1 − *W_j_*) *E*(1 – *W_k_*). In practice, in the models considered here, these correlations correspond to the entries of the variance-covariance matrix Σ. These biases naturally extend to the partial correlations this study focuses on, with the partial correlations corresponding to the entries of the precision matrix Ω = Σ^−1^.

Zero-inflation has a second effect on the inference of correlations. Even if one were able to perfectly identify the observations that emanate from the distribution of *N*, zero-inflation reduces the number of such observations and therefore the level of information available to infer the correlation structure of *N*. Therefore, even in a model that would perfectly represent the zero-inflation in a dataset, a substantial number of observations may be required to correctly infer correlations, be they marginal or partial. In the simulation study, we analyze whether increasing the number of observations suffices to compensate for this lack of information in zero-inflated models in the context of partial-correlation network inference.

### 2.2 The ZIPLN-network model

The ZIPLN-network model combines the properties of two pre-existing models, the PLN-network [Chiquet et al., 2019] and the ZIPLN model [Batardière et al., 2025].

Considering counts observations of *p* species in *n* samples, with *Y_ij_* the number of individual counts of species *j* found in sample *i*, so that *Y_i_* is a p-dimensional vector of integers, the Poisson log-normal (PLN) model [Aitchison and Ho, 1989; Chiquet et al., 2021] models observations *Y* using a Gaussian latent variable *Z* with:

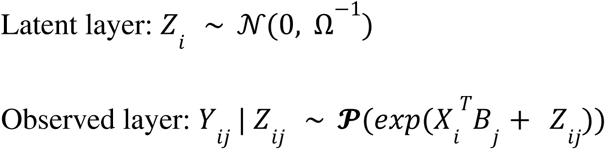

*X_i_* denotes the covariates used for sample *i*, *B_j_* the corresponding regression coefficients for species *j*. Just as in a regular generalised linear model, the *X_i_*^*T*^*B_j_* term accounts for abiotic effects on the abundances of species. We underline that the Poisson log-normal distribution is over-dispersed compared to a simple Poisson distribution.

To reduce the number of parameters to infer, the PLN-network models a sparse precision matrix Ω, in the graphical models tradition [Lauritzen, 1996]. By doing so, it allows the modeling of a sparse partial-correlation network that also corresponds to latent conditional dependencies. Indeed, given that *Z_i_* follows a Gaussian distribution, with variance-covariance matrix Ω^−1^, *Z*_−*j*_ and *Z*_−*k*_ (the *j* − *th* and *k* − *th* columns of *Z*) are independent *conditionally to all the other Z*_−*l*_ if and only if Ω*_jk_* = 0 [Whittaker, 2009]. It ensues that Ω gives an association network of partial correlations between species. Modeling the association network in a latent multivariate Gaussian layer is common to several JSDMs (see section 2.3), owing to the lack of a satisfying approach to model statistical dependencies in multivariate Poisson distributions [Besag, 1974, Chiquet et al., 2019]. In the observed layer, one has: *Cov*[*Y_ij_Y_ik_*] = *E*[*Y_ij_*]·*E*[*Y_ik_*]·(*exp*(Σ*_jk_*) − 1) so that the sign of the observation-layer covariance follows that of the latent layer. While the latent layer’s correlation structure cannot be directly transposed to the observation layer, the partial correlation structure of the latent layer remains informative, as it removes the effects of mediator species [Popovic et al., 2019]. The specificity of the PLN-network inference procedure [Chiquet et al, 2019] consists in adding an *l*_1_ penalty on the off-diagonal values of Ω to obtain a sparse partial-correlation network and select the most meaningful correlations. A LASSO-like approach adapted to graphs, called graphical-LASSO [Friedman et al., 2008], is used to this end.

Selecting the *l*_1_**-**penalty to apply can be done using different statistical criteria such as the AIC or BIC. In our study, we used the StARS method (Stability Approach to Regularisation Selection) [Liu et al., 2010]. This approach resamples a subset of the data multiple times, computes a new association network for each subsample and allows one to select a penalty that removes the associations that are present in less than x % of the networks, this proportion being fixed at 80 % in our study.

So far, we have described a model that assumes that all observations arise from a Poisson distribution (with an overdispersion arising from the latent variable). This means that any 0 observed in the data is explained by the effects contained in *X* or the correlations with other species through latent correlations. However, a particular species might be absent from a sample for structural reasons, not related to its rarity or abiotic effects, such as the existence of range limits.

The ZIPLN (Zero-inflated PLN) model [Batardière et al., 2025] aims at addressing this question. Instead of presuming that the data are produced by a Poisson distribution only, it assumes a mixture of a Dirac delta-distribution centered at zero to model zero-inflation and a Poisson distribution to model counts. To this end, it uses an additional binary latent variable *W* that follows a Bernoulli distribution. If *W_ij_* = 1, *Y_ij_* is automatically equal to 0 and if *W_ij_* = 0, *Y_ij_* is drawn from a Poisson distribution as with the PLN-network approach:

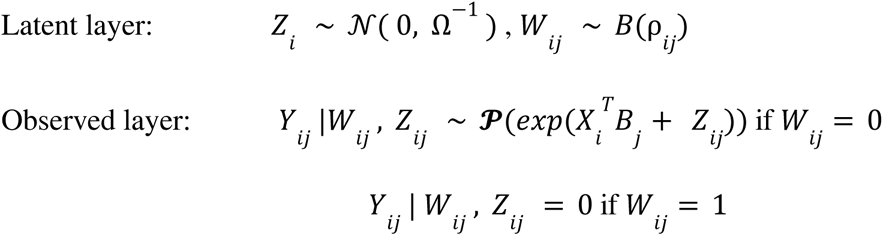

The parameters of W’s distribution can depend on the species only (ρ*_ij_* = ρ*_j_*), the sample only (ρ*_ij_* = ρ*_i_*), or the covariates, through a linear effect, with species-specific regression coefficient. In the latter case, we define 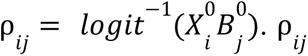 then depends on both *i* and *j*, but through covariates so that this does not lead to an overparametrization of the model. This means that it is possible to account for the fact that some species are more likely to be absent from a sample than others, but also that the probability of having a zero for a given species can depend on specific covariates.

Batardière et al. (2025) do not include the penalized approach for the inference of Ω in their model. The novelty of the ZIPLN-network model consists in mixing the ZIPLN and PLN-network approaches by modeling zero-inflation as is done in the ZIPLN model while also penalizing Ω off-diagonal values.

The ZIPLN-network model relies on variational Expectation-Maximization for parameter inference [Dempster et al., 1977, Wainwright and Jordan, 2008]. Its implementation is included in the publicly available R package PLNmodels [https://cran.r-project.org/package=PLNmodels] - a package that also implements the other PLN models [Chiquet et al., 2021].

In terms of notations, a PLN-network model run with *d* covariates named *X*_1_ to *X_d_* with an intercept to analyze species distribution matrix *Y* is denoted PLN-network(Y ∼ 1 + *X*_1_ + … + *X_d_*). The “1” term corresponds to the intercept, meaning that the species mean abundance contains a species-specific term.

Similarly, a ZIPLN-network model run with covariates *X*_1_ to *X_d_* is denoted

- ZIPLN-network(Y ∼ 1 + *X*_1_ + … + *X_d_* | zi = species) for a species-dependent zero-inflation,
- ZIPLN-network(Y ∼ 1 + *X*_1_ + … + *X_d_* | zi = site) for a site-dependent zero-inflation,
- ZIPLN-network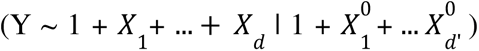 for a zero-inflation depending on an intercept and covariates 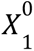 to 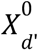.

Guidance on getting started with the ZIPLN-network model using R package **PLNmodels** is provided in Appendix S6.

In terms of interpretation, the network returned by the model is the precision matrix of the latent Gaussian layer of the Poisson component. Its entries are therefore residual partial correlations, computed once the effects of the environmental covariates have been removed and, in a zero-inflated model, after the effects attributed to the zero-inflation component have also been removed.

The practical consequence is that the two models weight shared zeros differently. Where a non-zero-inflated JSDM reads a pair of species jointly absent from the same sites as evidence of a conditional dependency, a zero-inflated JSDM may assign those zeros to its zero-inflation component, in which case the association weakens or disappears; conversely, a pair whose zeros are absorbed by that component may emerge with a stronger association. This does not by itself establish which of the two networks is closer to the truth, since shared zero patterns may equally reflect environmental effects or genuine ecological relationships. What it does provide is a way to single out the associations that rest on shared zeros, and hence to bring ecological knowledge to bear when interpreting them.

The ZIPLN-network model and the proposed inference strategy also have several limitations. First, the variational inference strategy [Chiquet et al., 2019] makes the derivation of confidence intervals for the estimators difficult. Although some work has been done in this direction [Batardière, 2024] the model does not provide uncertainty estimates. It also lacks the inclusion of random effects that would allow for the modeling of shared effects in community data, as other JSDMs propose [Brooks et al., 2017; Niku et al., 2019; Ovaskainen and Abrego, 2020].

In the simulation study, three approaches from the PLN models family are included. These are the ZIPLN-network model - that is the focus of our study - and the two models it emerges from: the PLN-network and (unpenalized) ZIPLN models. Studying the results of the PLN-network model allows us to analyze the effects of zero-inflation on partial-correlation network inference for a model that does not account for it.

A further important point is that the ZIPLN model does not apply a penalty to the precision matrix. This allows us to see how critical the focus on the modeling of a sparse precision matrix is when testing the inference of partial-correlation networks. To make its results more comparable with those of the reference models (see section 2.3), we do not directly extract the precision matrix it infers but rather use the full-rank variance-covariance matrix and apply the Graphical-Lasso algorithm to obtain a sparse precision matrix (as is done with the Hmsc and gllvm reference models, see section 2.3).

### 2.3 Reference JSDMs

We compared the ZIPLN-network results to those obtained with JSDMs implemented in two packages: Hmsc [Ovaskainen et al., 2017, Ovaskainen and Abrego, 2020, Tikhonov et al., 2020] and gllvm [Niku et al., 2019]. These packages allow the use of different distributions to model observations, including Poisson for count data and Bernoulli for presence-absence. For the sake of comparability, we focus on the zero-inflated Poisson distribution-based models.

We now describe the gllvm and Hmsc models. Let us first consider the models with no zero-inflation modeling. In such cases, when using a Poisson distribution, the modeling framework is essentially the same in PLN-network, gllvm and Hmsc, with the model linking observations to explanatory covariates and a latent Gaussian variable through a log-link function.

The main difference lies in the fact that, while the (ZI)PLN-network model penalizes the precision matrix to reduce the number of parameters, the gllvm and Hmsc models resort to a low-rank approximation of the variance-covariance matrix, that is, the two families do not natively target the same object (see Introduction).

The precision matrix based and variance-covariance matrix based can both be of interest and there is no general consensus about using one or the other. This paper focuses on the precision matrix and the associated partial-correlation network, as is often done in statistics in the graphical models literature [Lauritzen, 1996, Whittaker, 2009] and in other fields such as microbial studies [Kurtz et al., 2015].

One could argue that it suffices to invert the variance-covariance matrix that Hmsc and gllvm infer to obtain the precision matrix, thereby considering the same framework as the PLN-network does. However, the Hmsc and gllvm use a low-rank approximation of the variance matrix (with their rank being determined by the number of latent factors) to reduce the number of parameters, making them singular.

To address this, one option would be to run these models with as many latent factors as there are species. However, in this case, that was not efficient: not only did the quality of the partial-correlation networks inference not improve, but also, in the case of the gllvm model, the running time increased tremendously while convergence issues arose (Appendix S2).

A second option is making the inferred variance matrices full-rank by adding a small offset on their diagonal. However, this can strongly distort the corresponding inverted matrix. Alternatively, and the option we used here, as in practice it appears to work better, is to apply a Graphical-Lasso algorithm [Friedman et al., 2008] to the gllvm or Hmsc inferred variance matrix. The algorithm works even if the matrix is singular. Another advantage of this approach is that one can control the level of sparsity on the precision matrix with the Graphical-Lasso penalty. This offers a natural way to select the most statistically significant associations.

Finding a way to infer precision matrices from each model is important because our assessment of a method’s ability to retrieve the correct partial-correlation network is based on the rate of true *versus* false positives (that is true and false associations) at a given level of sparsity / confidence.

Another difference between the (ZI)PLN-network model, and the gllvm and Hmsc models, is that the latter additionally allow for the inclusion of species traits effects (both models) and of spatial and phylogenetic effects (Hmsc only) - features that are not part of the present study.

There is an option with gllvm to use a zero-inflated Poisson distribution to address zero-inflation - an approach comparable to the solution provided in the ZIPLN-network. Hmsc, in contrast, offers no zero-inflation implementation as it appears more difficult to incorporate in its hierarchical framework. The Hmsc model can however be used as a hurdle model where presence-absence is modeled with a Bernoulli distribution, and abundance modeled with a Poisson distribution, conditionally on presence. In practice, both steps are done independently and we only need to run the second one to obtain the desired association network.

The Hmsc estimation procedure resorts to Bayesian inference whereas gllvm uses a frequentist approach. Both gllvm and Hmsc are implemented in R packages (see Niku et al. (2020) for gllvm and Tikhonov et al. (2020) for Hmsc) that were used for this study. For gllvm we used the default parameters (increasing the number of latent dimensions did not seem to improve the results). For Hmsc, we first checked the convergence of the Bayesian inference procedure on several examples and then proceeded to run the model with thin = 10, samples= 1000 and transient = 500.

#### A note on glmmTMB

The glmmTMB package [Brooks et al., 2017] implements generalized linear mixed models with a wide range of emission laws and zero-inflation structures, and inter-species dependence can be introduced through a random effect that is either low-rank or full-rank. With a zero-inflated Poisson distribution and a full-rank random effect, it specifies exactly the unpenalized ZIPLN model, with the same parameterization; only the fitting procedure differs, glmmTMB maximizes a Laplace approximation of the likelihood through TMB [Kristensen et al., 2016] where PLNmodels maximizes a variational lower bound. The generality of the Laplace approximation is the strength of the approach, as a single implementation covers model families that would each otherwise require a dedicated derivation. However, a full-rank random effect over *p* species carries *p*(*p* + 1) / 2 unstructured covariance parameters, and in our simulation settings the optimizer rarely reached convergence, at a computational cost roughly two orders of magnitude above that of the tailored variational procedure (in our settings, minutes against seconds per fit). We therefore did not include glmmTMB in the systematic comparison. We stress that this reflects the cost of a generic fitting procedure in this regime, and not a limitation of the model it specifies, since it is precisely the same as our own.

When it comes to the object of our analysis, partial-correlation network inference, the models included in the comparison vary in three aspects: the way they represent zero-inflation, their modeling of the partial correlation structure and the inference method they resort to. Table 1 summarizes the different approaches used in the simulation study, their models and the inference methods they resort to. While the simulation study (Section 3.1) tells us how each one performs, it does not allow one to perfectly tease apart the effect of each factor on these differences, although we can formulate several hypotheses based on the difference summarized in Table 1. This emerges as one of the limits of this study.

**Table 1:**
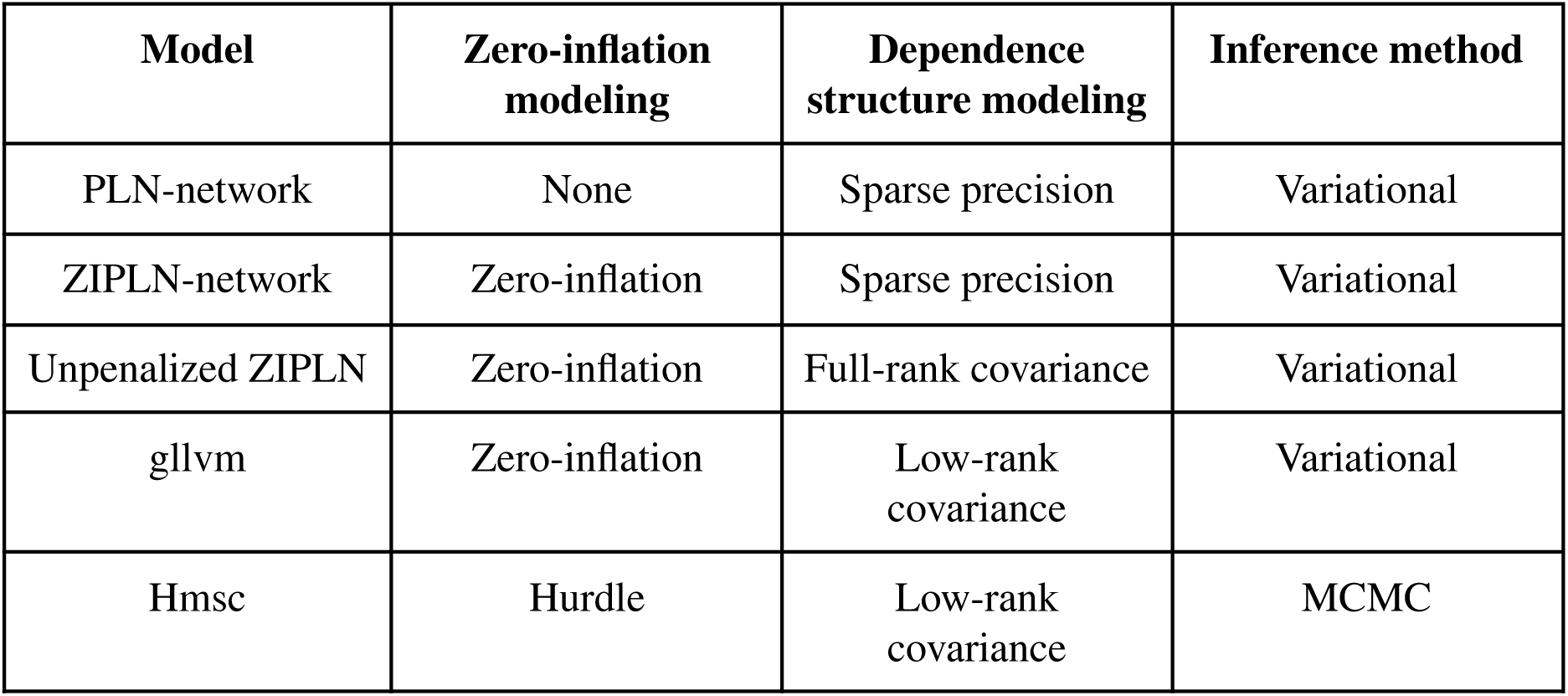
comparison of the JSDMs included in the study.

| **Model** | **Zero-inflation modeling** | **Dependence structure modeling** | **Inference method** |
| --- | --- | --- | --- |
| PLN-network | None | Sparse precision | Variational |
| ZIPLN-network | Zero-inflation | Sparse precision | Variational |
| Unpenalized ZIPLN | Zero-inflation | Full-rank covariance | Variational |
| gllvm | Zero-inflation | Low-rank covariance | Variational |
| Hmsc | Hurdle | Low-rank covariance | MCMC |

### 2.4 Simulation protocol

We compared the results of the ZIPLN-network model to those of the PLN-network, unpenalized ZIPLN, Hmsc and gllvm models on simulated data.

Our goal was to compare each model’s ability to retrieve partial correlation species association networks, to assess the computation time they require and to test three related predictions:

1. Failing to account for the zero-inflation in count-based community data impairs partial-correlation network inference.
2. Network inference is impaired in line with the increasing importance of zero inflation.
3. Increasing the number of observations can compensate for the lack of information induced by zero-inflation in a zero-inflated model.

To this end, we simulated abundance data with various types and levels of zero-inflation and compared the networks inferred by each model.

Sparse networks were generated using an Erdös-Rényi model (no particular structure), a preferential attachment structure (edges are attributed progressively with a probability proportional to the number of edges each node is already involved in), or a community structure (in the Stochastic Block Models sense [Holland et al., 1983]). This allowed us to compare the results given different network structures. Using the **igraph** package [Csardi and Nepusz, 2006], we generated an adjacency matrix *G* corresponding to each structure. Then a precision matrix Ω was created with the same sparsity pattern as *G*, as follows: Ω’ = *G* × *v*, Ω = Ω’ + *diag*(|*min*(*eig*(Ω’))| + *u*) with *u*, *v* > 0 two scalars. A higher *v* implies stronger correlations whereas a higher *u* implies a better conditioning of Ω.

Following [Chiquet et al., 2019], we fixed *v* = 0. 3, *u* = 0. 1 in the simulations. Each non-zero value of Ω had a probability 0.4 of being turned to its opposite so as to ensure that Ω contained negative values too.

We did not simulate data under the PLN model but instead, following Chiquet et al. (2021), used a compositional model so as to ensure fairer tests. We first drew a lognormal variable with *a_i_* ∼*LN*(*X_i_B*, Ω^-1^). These pseudo-abundances were turned into proportions with a logistic transformation 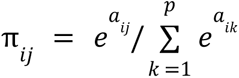. Finally each count vector *Y_i_* was drawn from a multinomial distribution using the vector of proportions given by π*_i_*.

To further test over-dispersion we also ran simulations under a Negative-Binomial distribution with and without zero-inflation. The results for this set of simulations are presented in Appendix S3.

The simulated data already contained a number of random zeros in the abundance data, produced by the compositional model. We then proceeded to incorporate zero-inflation, emulating structural zeros. We tested all three types of zero-inflation: species-dependent, site-dependent and covariate-dependent. For each type, we tested four levels of zero-inflation. For species or site dependent zero-inflation, the reference zero-inflation levels correspond to the species (or sites) being split evenly into 4 groups, with respectively 10, 25, 50 and 75 % of zero-inflation. Two lower levels were tested, corresponding to respectively 10 and 50 % of this reference level, as well as a level 10 % higher. For covariate-dependent zero-inflation, a single categorical variable was used, with 3 possible values. The species were split into 5 groups depending on their response to this categorical variable. This results in 15 possible zero-inflation percentages, regularly spaced out between 10 % and 80 %, with, again, two lower levels of zero-inflation (25 and 50 % of the reference) and a higher level (10 % higher). The zero-inflation levels are denoted 1 to 4 on the Figures from lowest (few structural zeros) to highest (many structural zeros).

We ran simulations with *n* = 50, 100 or 300 sites and *p* = 25 or 50 species. We used an intercept and a single continuous covariate (denoted *X* in the model) drawn between 0 and 1. Each parameter configuration (that is each combination of *n*, *p*, network structure, zero-inflation type, zero-inflation level) was repeated 20 times.

For each method, we obtained networks (in this case, precision matrices) with different levels of sparsity either through penalization ((ZI)PLN-network) or thresholding (gllvm, Hmsc). The different methods could therefore be compared in terms of AUC (Area Under the receiver operating characteristic Curve) with the true positive rate (fraction of non-zero entries in the simulated precision matrix that are identified) and the false positive rate (fraction of the non-zero entries in the inferred precision matrix that do not correspond to a true association in the simulation) computed for each sparsity level.

The AUC gives a general idea of each model’s ability to correctly retrieve the association network but it does not assess the precision of point estimates. Although our focus here is on the retrieval of a network of associations and hence on the distinction between zero and non-zero values in the precision matrix, we also consider the RMSE (root mean squared error) between the inferred and real partial correlation matrices (obtained after normalising the precision matrices). We consider the absolute RMSE and the relative difference between the RMSE and that obtained with an empty network (that is, with all partial correlations set at 0) to ensure that good RMSE results are not a simple artifact owing to the sparsity of the network. For the (ZI)PLN-network models we computed the precision matrices obtained with penalties selected either with the BIC or with the StARS method (with levels 80 % and 90 %). Given that the BIC and StARS criteria give access to a sound method to select a specific sparsity level, we also computed a point-wise F1-score for each precision matrix. Finally, we measured the computation time required by each method in each configuration.

We additionally ran simulations under sparse variance-covariance matrices and tested each model’s ability to retrieve the support of the variance-covariance matrix in this context with the AUC. In this context, the Hmsc AUC was computed using the posterior probabilities the model outputs, to select non-zero-values, whereas for the gllvm, PLN and ZIPLN we used a simple thresholding. This allowed us to test whether there were important empirical differences in the influence of zero-inflation on the models’ abilities to retrieve the support of the variance *versus* precision matrix and whether the models compared differently in this context. The results of this analysis are presented in Appendix S1.

Code for the simulation study can be found in the git repository ZI_JSDM_for_association_network_inference_from_ZI_count_based_community_data

### 2.5 Tropical freshwater fish data

To illustrate how the ZIPLN-network approach can be used to study species community data, we used a freshwater fish survey from Trinidad. Trinidad is an island in the southern Caribbean, located 12 km from Venezuela. It is home to a rich freshwater fish fauna of both Antillean and South American origin that has been extensively investigated [Phillip et al., 2013].

The data set used here is described in Magurran et al. (2018). The original fieldwork was undertaken as part of ERC AdG BioTIME (250189), and received ethical clearance from the ERC. The survey was carried out in the Northern Range of Trinidad, a mountain system in the north of Trinidad where a series of parallel rivers are found. The fish were surveyed at 16 sites in eight Northern Range rivers over a 5-year period from December 2010 to August 2015. In each river, two sites were surveyed, one considered as “disturbed” due to human recreational activities, while the other was “undisturbed” [Magurran et al., 2018]. Each site was surveyed four times a year, at the beginning and end of both the dry (January to May) and wet (June to December) season [Deacon et al., 2015, Magurran et al., 2018]; the first sample was removed due to differences in sampling effort [Magurran et al., 2018]. Hence, 19 surveys are considered for each site. The dataset contains 20 species, after filtering out those that had been observed less than twice in the dataset. For the sake of readability, we use shortened versions of the species names in the figures; their complete names can be found in Appendix S4.

The dataset contains numerous zeros (Figure 1). We assumed that it does not contain false zeros, given the rigorous sampling protocol used [Magurran et al., 2018] (while false zeros may remain, there would be no way to identify them). Some of the zeros may arise from sampling variability, owing to the overall rarity of several species, like *Synbranchus marmoratus* or *Awaous banana*. Rarity-related zeros should mostly be represented in the mean effect of the Poisson distribution in the ZIPLN-network model, through the intercept. Some species, while relatively abundant in many sites, appear not to be found at all in several streams (Figure 1). For example *Andinoacara pulcher* is completely absent from the upper Aripo stream, and *Ancistrus maracasae* is not found in the Quare and Turure streams. These zeros seem to be structural zeros as defined by Blasco-Moreno et al. (2019). They are likely owing to natural limits in the dispersal of the species. Indeed, in the context of these data, natural barriers, like waterfalls, can prevent a species from reaching a given site even if the abiotic and biotic contexts could have suited it. These structural explanations for the presence of zeros in the dataset make it particularly suitable to illustrate the use of the ZIPLN-network model.

**Figure 1:**
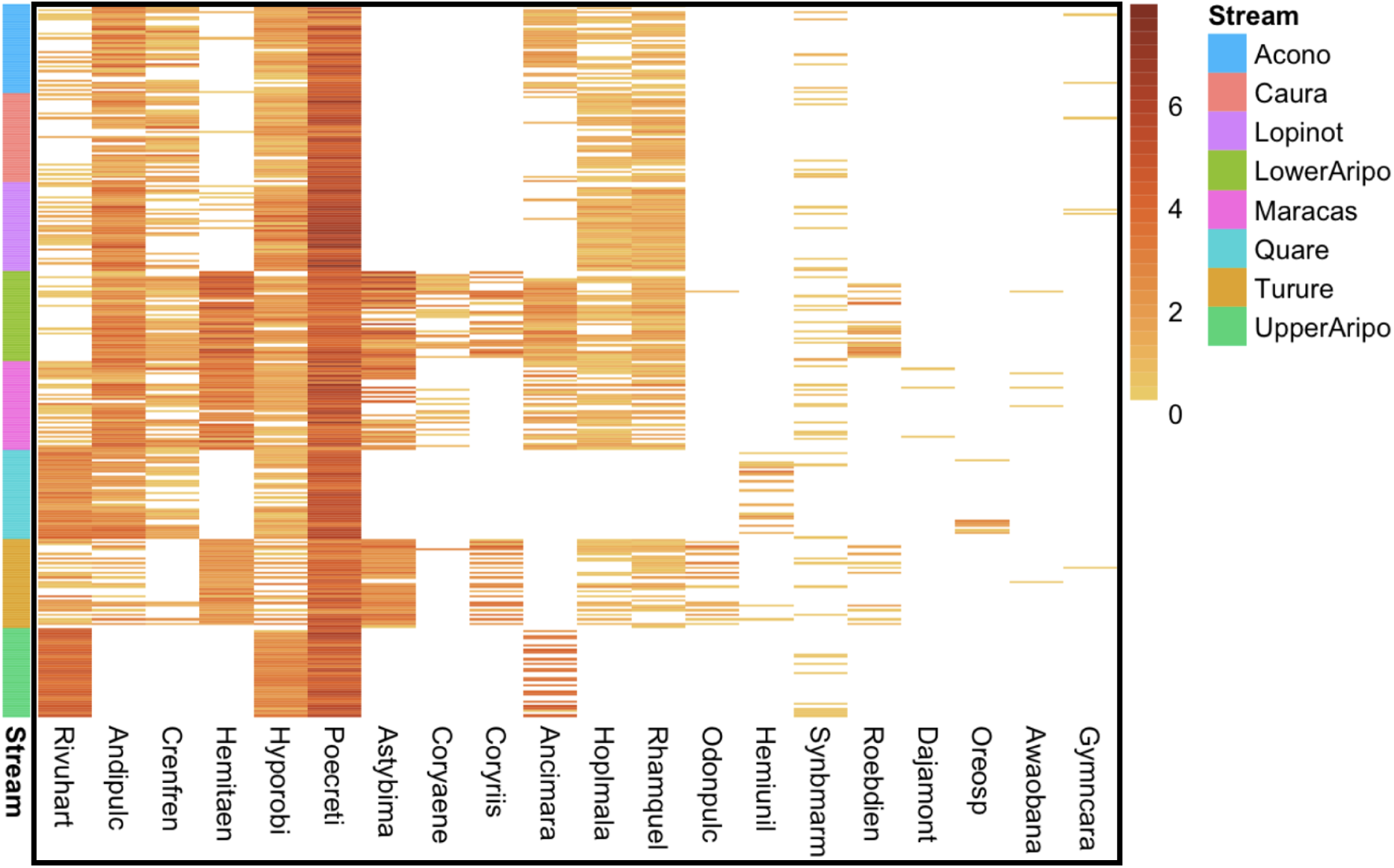
Species log-abundances (log(1 + species count), in the dataset, sorted by stream. Blank cells indicate species absences. Each column corresponds to a species, each row corresponds to a combination (site, time step). The rows are grouped by stream (there are two sites per stream).

We hypothesized that accounting for the zero-inflation in the analysis of this data set would remove several potentially spurious associations. In principle it also supports a more precise analysis as it can help separate different types of zeros. We additionally hypothesized that some associations owing to the presence of structural zeros may appear in the PLN-inferred networks but not in the ZIPLN-inferred ones.

### 2.6 Choice of covariates

To account for abiotic effects, we ran a principal component analysis (PCA) on a set of descriptive variables. The variables used were: width, depth and volume of the stream, altitude, water conductivity, dissolved oxygen concentration, pH, temperature, and turbidity, as well as several quantitative descriptors of the substrates found in the river and a visual estimate of the canopy cover. We only kept the first two principal components, denoted *PC1* and *PC2*; using more did not significantly improve or modify the results. *PC1* mostly corresponded to water descriptors (depth, volume, temperature) whereas *PC2* corresponded to descriptors of the substrates in the river (see Appendix S5). Including these covariates avoids finding statistical associations owing to their joint effect on two species.

We additionally accounted for two categorical variables: disturbance (binary variable, i.e., whether the site was *a priori* labeled as disturbed) and river identity. Together, these two variables characterise each geographical site. Accounting for the stream ID in this way allowed us to include a dispersal-related effect in the model: if a species was never found in a particular stream or in nearby sites, the model adjusted the coefficient of B corresponding to that stream, without attributing such an effect to the other abiotic covariates.

While it could also be of interest to consider the river descriptor as a random effect and thereby include inter-river correlations in the model, this is not permitted by the model at the moment.

For the PLN-network model, we assumed that all the covariates we included influence the abundances, whereas for the ZIPLN-network model, we included the stream effect in the zero-inflation part of the model only, and used PC1, PC2, and the disturbance variable for the abundance.

In total, we studied the species association networks obtained from six different models (three PLN-network and three ZIPLN-network models):

PLN-network(Y ∼ 1 + PC1 + PC2)
PLN-network(Y ∼ 1 + PC1 + PC2 + disturbance)
PLN-network(Y ∼ 1 + PC1 + PC2 + disturbance + stream)

ZIPLN-network(Y ∼ 1 + PC1 + PC2 | zi = species)
ZIPLN-network(Y ∼ 1 + PC1 + PC2 + disturbance | zi = species)
ZIPLN-network(Y∼ 1 + PC1 + PC2 + disturbance | 1 + stream)

## 3. Results

### 3.1 Simulation study

As expected, an increasing level of zero-inflation appears to make it impossible for a non-zero-inflated model like the PLN-network model to correctly retrieve a network inference (Figure 2), with all the different network configurations (Figure 3). Within the range of sample sizes tested (up to six times the number of species), increasing the number of observations does not allow the PLN-network model to recover from zero-inflation (Figure 4). The network structure also appears to have an impact on the result, although a limited one, with the preferential attachment network being easier to retrieve than the other two for the ZIPLN and ZIPLN-network models when the zero-inflation is species-dependent (Figure 3).

**Figure 2:**
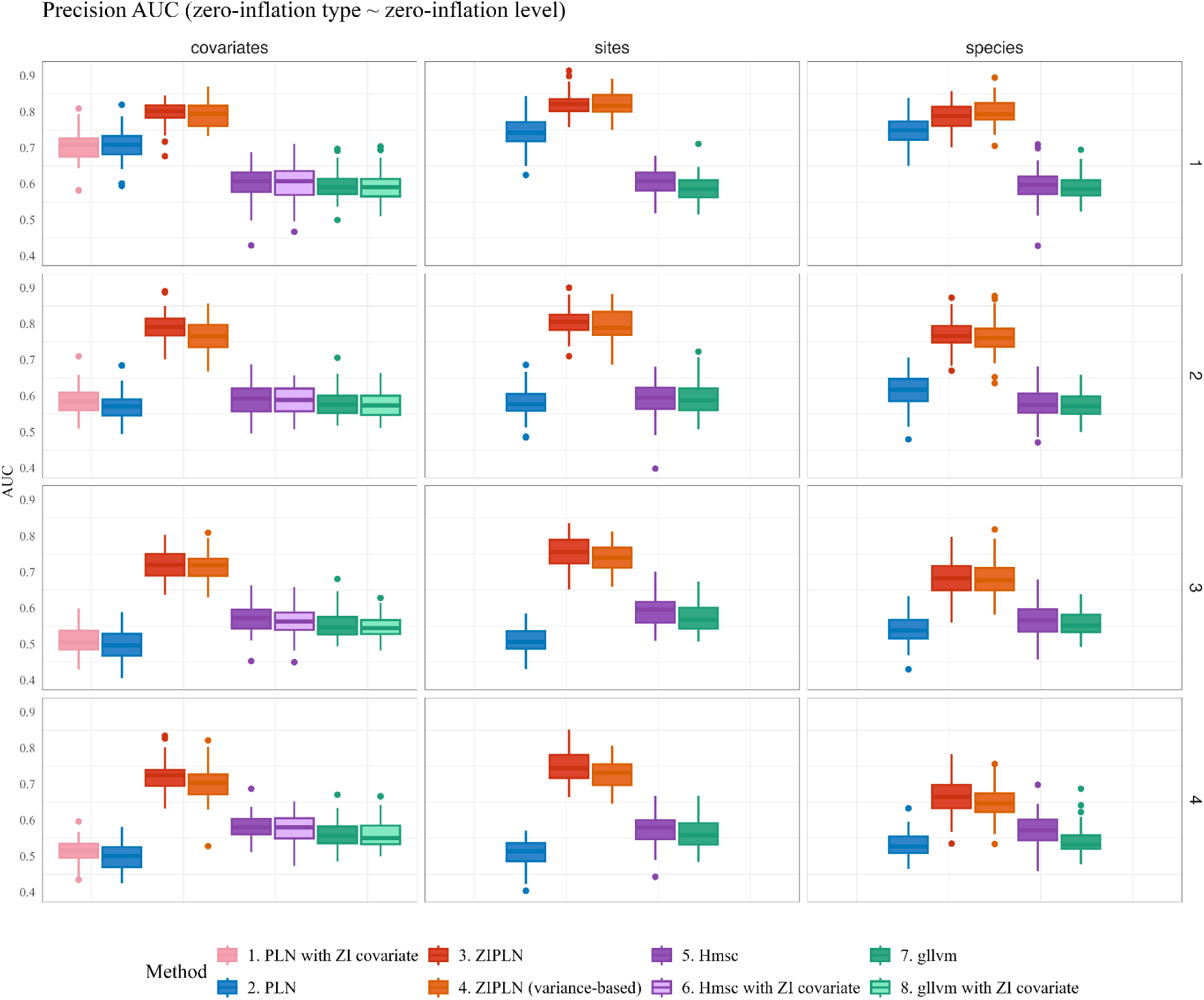
Boxplot of the AUCs obtained with the different network inference methods tested, as the zero-inflation type and zero-inflation level (1 = lowest, 4 = highest) vary, with *n* = 100, *p* = 50. The AUCs obtained by the ZIPLN model are consistently higher than those obtained with the other models; this difference is more striking when the zero-inflation is covariate or sites-dependent. Each boxplot corresponds to 60 replicates (20 for each of the three tested network structures).

**Figure 3:**
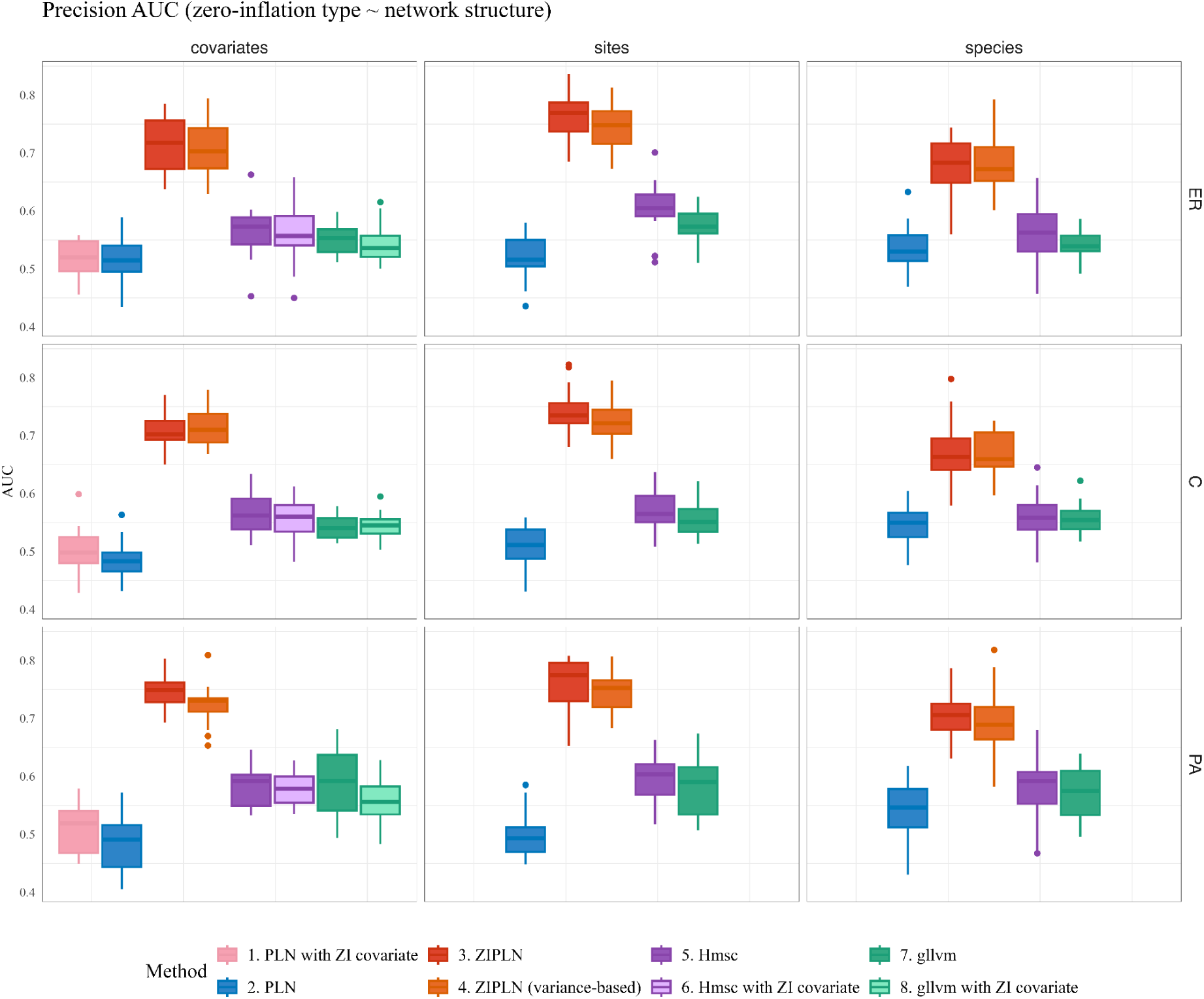
Boxplot of the AUCs obtained with the different network inference methods tested as the network structure and zero-inflation type vary for a zero-inflation fixed at the reference level (level 3 on the other plots), *n* = 100, *p* = 50 (**ER:** Erdös-Rényi, **PA:** Preferential Attachment, **C:** Community). Each boxplot corresponds to 20 replicates.

**Figure 4:**
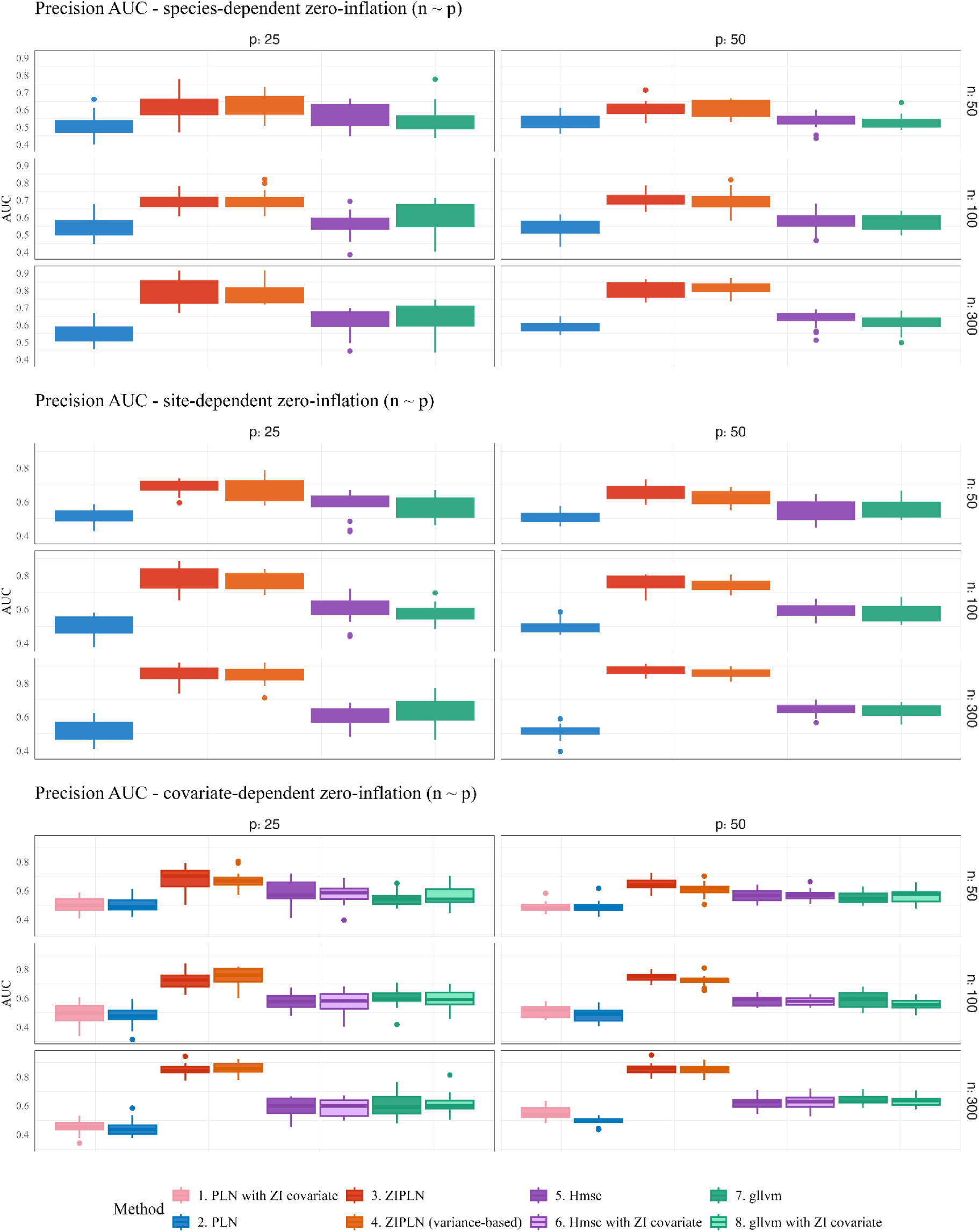
Boxplot of the AUCs obtained with the different network inference methods tested as the number of sites (*n*) and of species (*p*) vary for the different zero-inflation types, with zero-inflation levels fixed at the reference level (level 3 on the other plots). Each boxplot corresponds to 60 replicates (20 for each of the three tested network structures).

The AUCs indicate that the zero-inflation modeling by the ZIPLN-network model efficiently allows for the retrieval of the partial-correlation network (Figures 2 to 4). Interestingly, it appears that a partial-correlation network resulting from a penalized model that directly outputs a sparse precision matrix does not outperform that of the network obtained by inverting the full-rank precision matrix of a non-penalized model.

Increasing the number of observations (Figure 4) with a zero-inflation at the reference level (level 3 on the figures) allows the model to get AUCs comparable to those obtained with the lowest level of zero-inflation observed on Figure 2, indicating the importance of the role played by the degradation of the information in a zero-inflated context.

While the gllvm and Hmsc models consistently obtain better-than-random AUCs, they delivered poorer results than the ZIPLN-network model (Figures 2 to 4). The variance-based results of the unpenalized ZIPLN model shows that this is not due to their primarily modeling marginal correlations, but may be linked to the use of a low-rank approximation. Increasing the ranks of the approximation did not prove efficient (Appendix S2).

When it comes to the results obtained for the variance matrix, Appendix S1 (simulations in a sparse variance context) shows that the marginal correlation structure is harder to retrieve as all the models obtain lower AUCs. The ZIPLN-network model appears to be able to obtain reasonable AUCs when the number of observations is great enough (300 observations for 50 species).

A clearer empirical sense of why zero-inflation can hinder the retrieval of partial correlations, emerges from the PLN-network and ZIPLN-network results on a specific simulation as is illustrated by Figure 5. Figure 5.C shows that the PLN-network model infers positive associations that are absent from the ground-truth network (Figure 5.A) between species 4, 5 and 6. We note that these species share a similar zero-inflation pattern (see column 4 to 6 on the heatmap of Figure 5.B). When the zero-inflation covariate is accounted for as a mean effect in the PLN-network model (Figure 5.D), only the association between species 4 and 5 remains, but numerous other spurious associations persist that are filtered out by the ZIPLN-network model (Figure 5.E).

**Figure 5:**
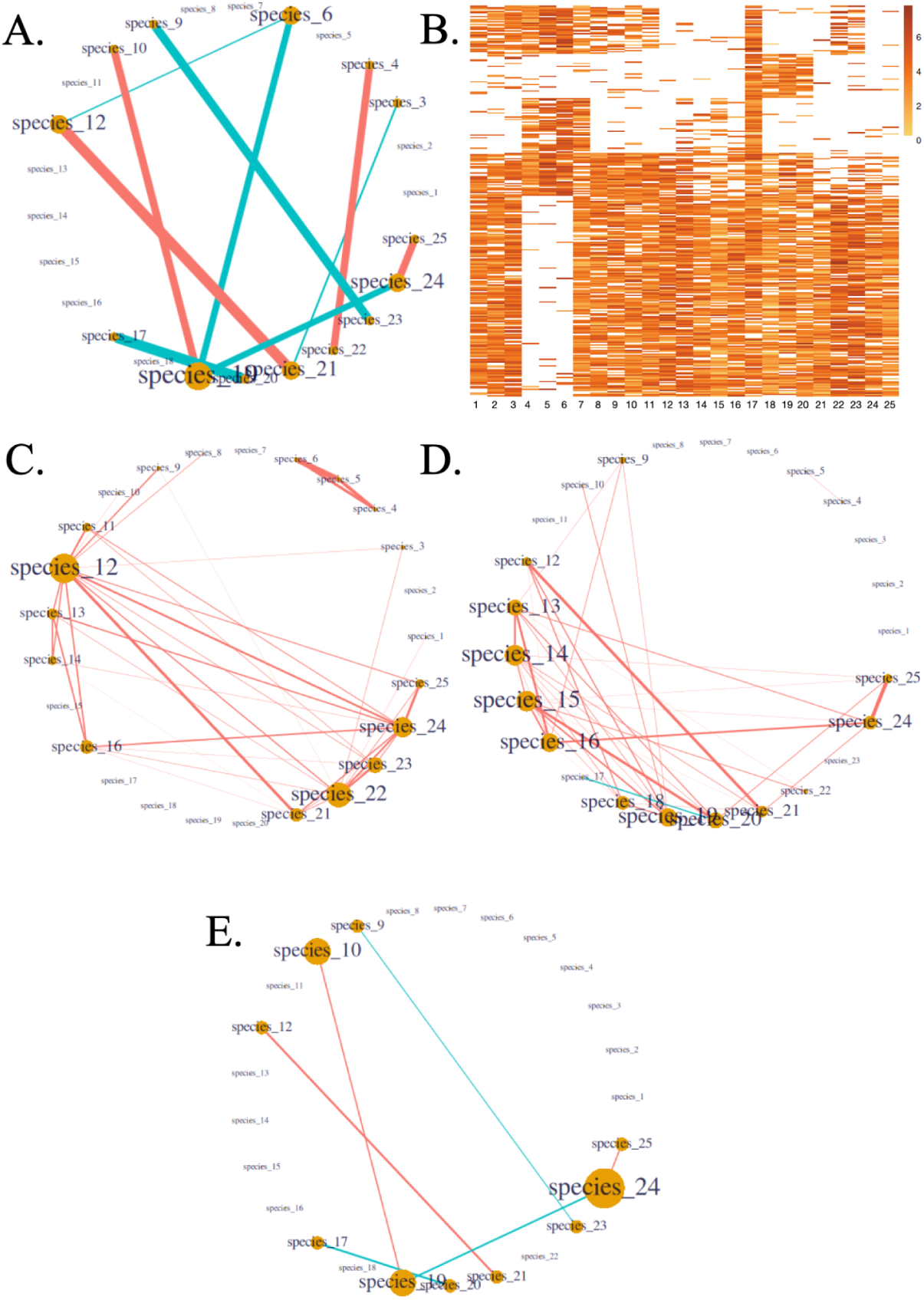
Illustrating how an association network is retrieved from zero-inflated abundance data with and without accounting for zero-inflation. Red edges correspond to positive associations and blue edges to negative ones.. **A.** Ground-truth network used for the simulation. **B.** Heatmap of the log-abundance generated for this simulation, with covariate-dependent zero-inflation, zero-inflation level being at the reference level (level 3 on Figure 2). **C.** Species association network inferred by the PLN-network model. **D.** Species association network inferred by the PLN-network model with the zero-inflation covariate included in the mean of the model. **E.** Species association network inferred by the ZIPLN-network model.

In terms of point-wise estimates of the network, an increasing zero-inflation significantly degrades the F1-score obtained by the PLN-network and ZIPLN-network model, with the ZIPLN-network being able to keep much higher F1-scores than the PLN-network model, as could be expected (Figure 6). These scores however, tend to remain relatively low (below 0.5 for zero-inflation levels 3 and 4), but increasing the number of observations significantly improves them (Figure 7). It also appears that the StARS method is better suited to choosing a penalty than the BIC. Changing the StARS threshold from 0.8 to 0.9 only appears to have a slight impact on the results.

**Figure 6:**
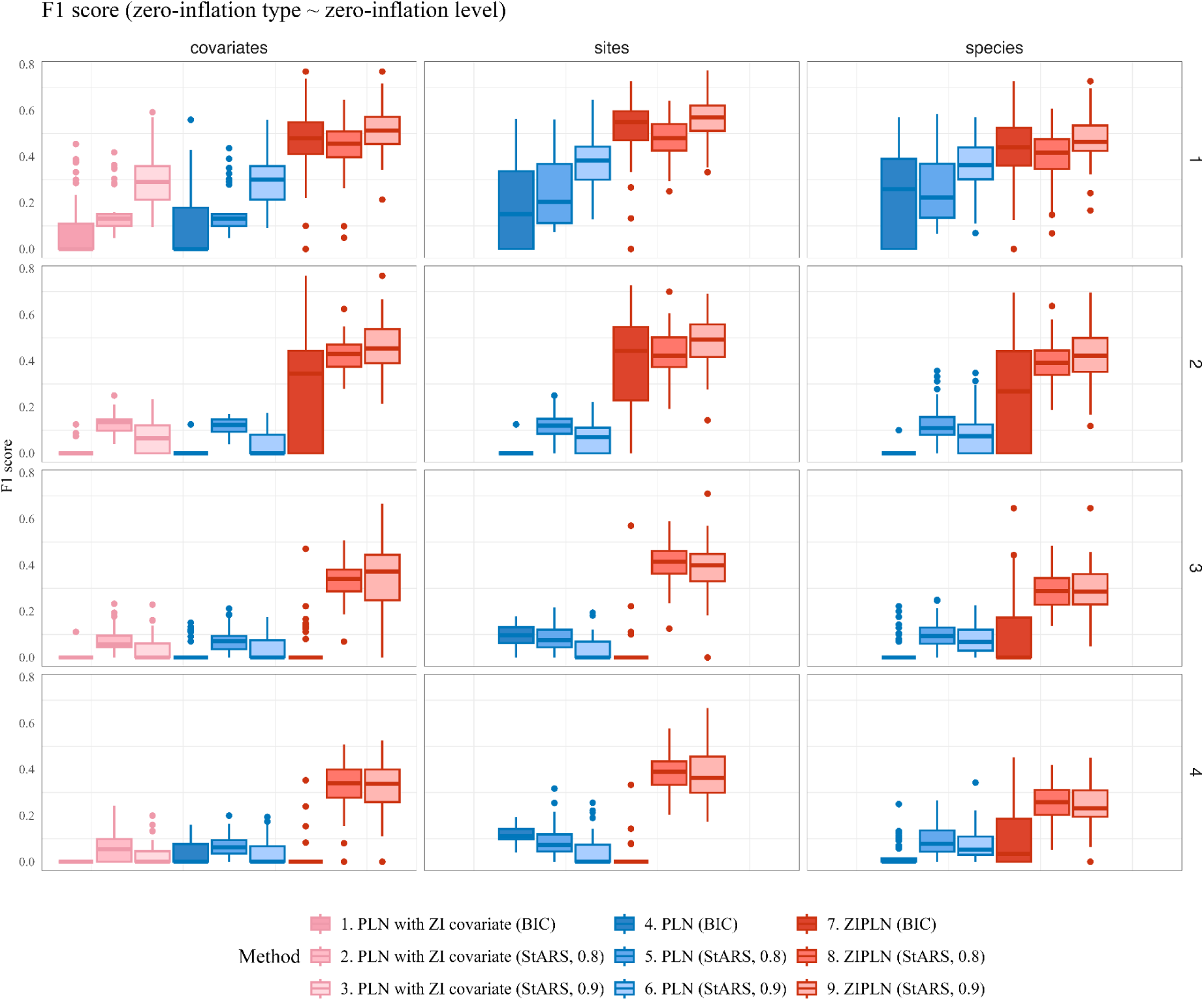
Boxplot of the F1 scores obtained with the penalized PLN and ZIPLN methods for different zero-inflation types and levels; for penalties selected with different criteria (either BIC or StARS at levels 0.8 or 0.9), with *n* = 100, *p* = 50. Each boxplot corresponds to 60 replicates (20 for each of the three tested network structures). Zero-inflation levels: 1 = lowest, 4 = highest.

**Figure 7:**
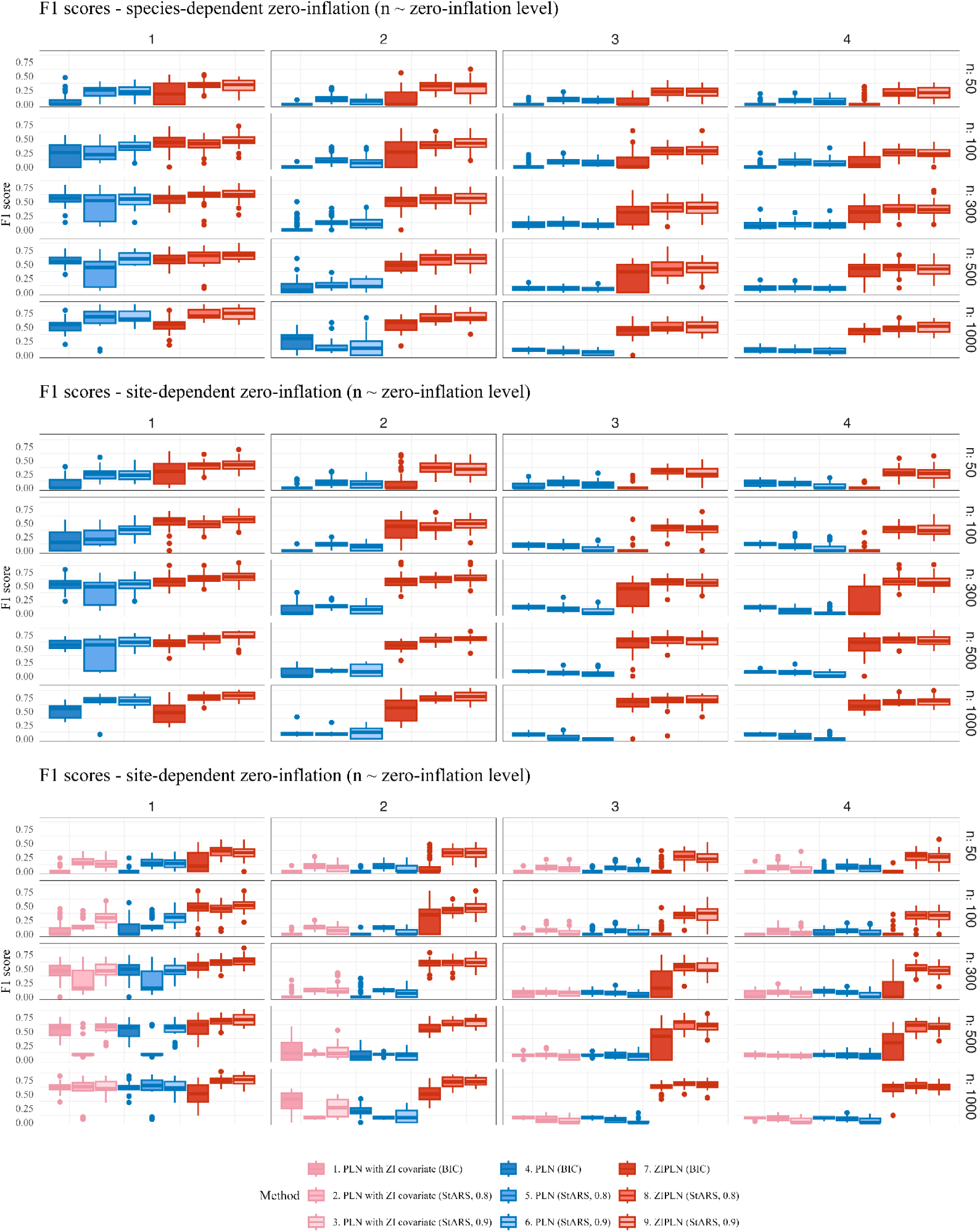
Boxplot of the F1 scores obtained with the penalized PLN and ZIPLN methods for penalties selected with different criteria (BIC or StARS at levels 0.8 or 0.9), as the zero-inflation level and the number of sites (*n*) vary for the different zero-inflation types. Each boxplot corresponds to 20 replicates. Zero-inflation levels: 1 = lowest, 4 = highest.

The partial correlation RMSE (Figures 8 and 9) gives another view of the point-wise estimated results as they do not, strictly speaking, assess the retrieval of the network (the non-zero values in the precision) but the distance to its values. An increasing zero-inflation seems to impact the RMSEs less than the AUCs, and the models show comparable performance with the PLN-network obtaining the worst RMSEs and the ZIPLN-network getting the best ones (Figure 8). The results of the ZIPLN-network are likely helped by the fact that it outputs a sparse precision matrix. This is why it is important to not only consider these RMSEs *per se* but also their difference with those an empty, uninformative network would obtain (Figure 9). Again, the ZIPLN-network performs better, consistently obtaining better RMSEs than the empty network, when the PLN-network results are the worst, especially when the zero-inflation is species-dependent.

**Figure 8:**
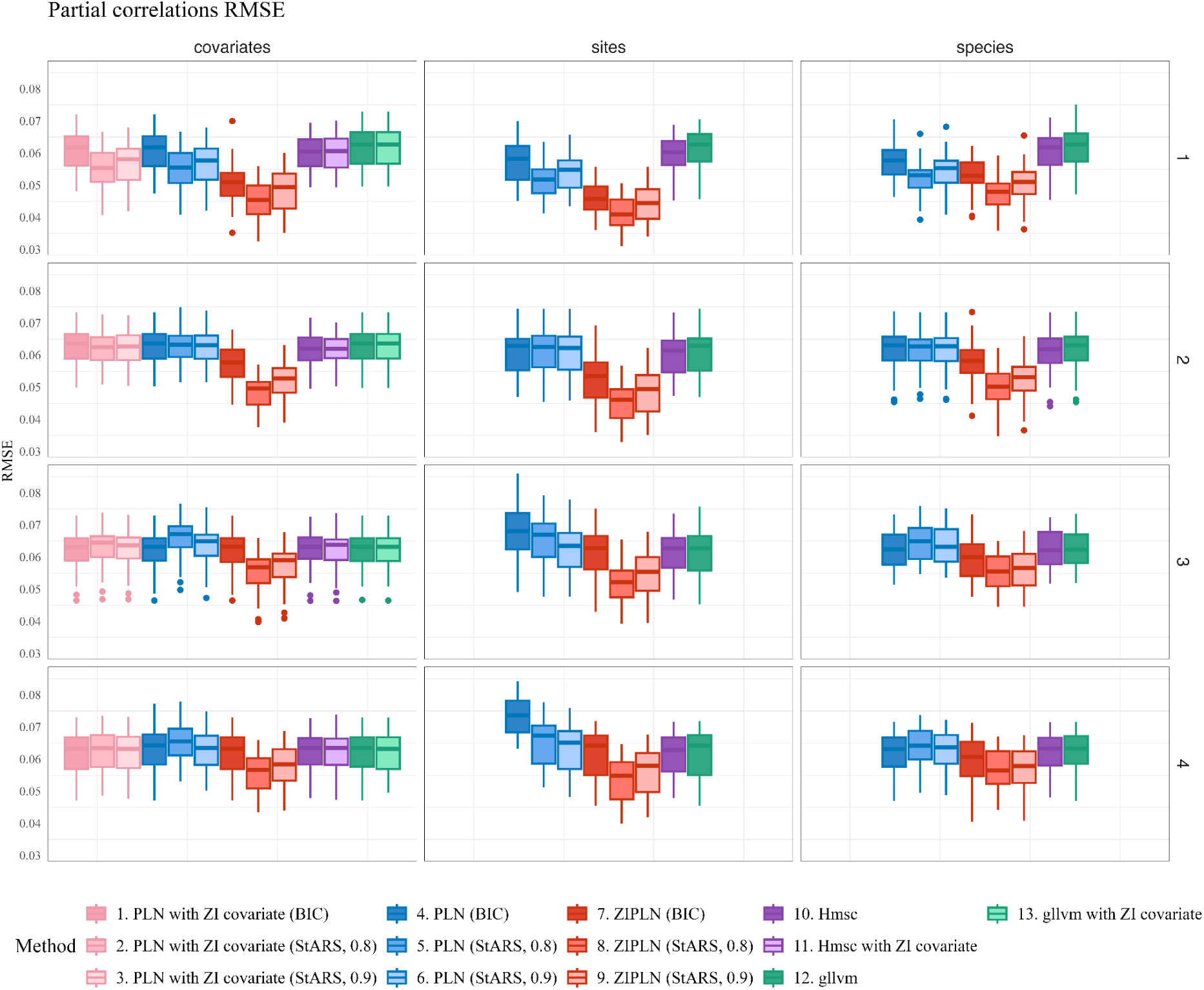
Boxplot of the RMSE obtained for the partial correlations by each method, with each criterion when applicable, for each zero-inflation configuration, with *n* = 100 and *p* = 50. Each boxplot corresponds to 60 replicates (20 for each of the three tested network structures). Zero-inflation levels: 1 = lowest, 4 = highest.

**Figure 9:**
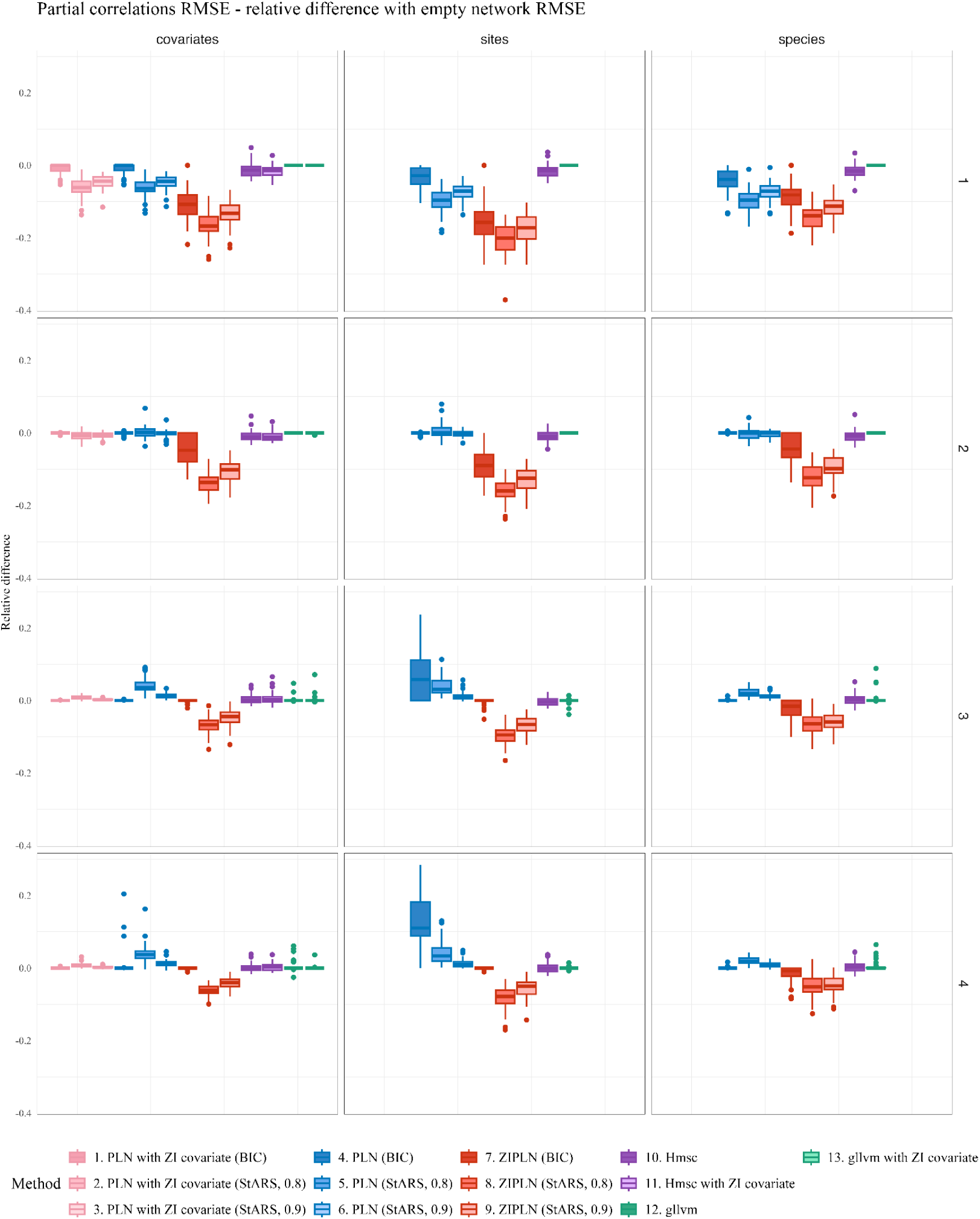
Relative difference between the partial correlations RMSE obtained by each method and each criterion (if applicable) and the RMSE obtained with an empty network (all partial correlations equal to 0), with *n* = 100, *p* = 50. Each boxplot corresponds to 60 replicates (20 for each of the three tested network structures). Zero-inflation levels: 1 = lowest, 4 = highest.

The same analysis can be done for the variance (Appendix S1, simulation study with a sparse variance matrix). It also shows that the marginal correlations are harder to estimate than the partial ones and that no model performs very well in this context.

Finally, the running times (Figure 10) required by the PLN-network, ZIPLN-network and gllvm models are comparable. Using the StARS criterion for (ZI)PLN-network is more time-consuming because it requires fitting the model multiple times to different subsamples of the data (the StARS cut-off level has no impact on the running time since it is only used after the multiple fitting). The Hmsc model however takes much longer owing to the Bayesian inference procedure.

**Figure 10:**
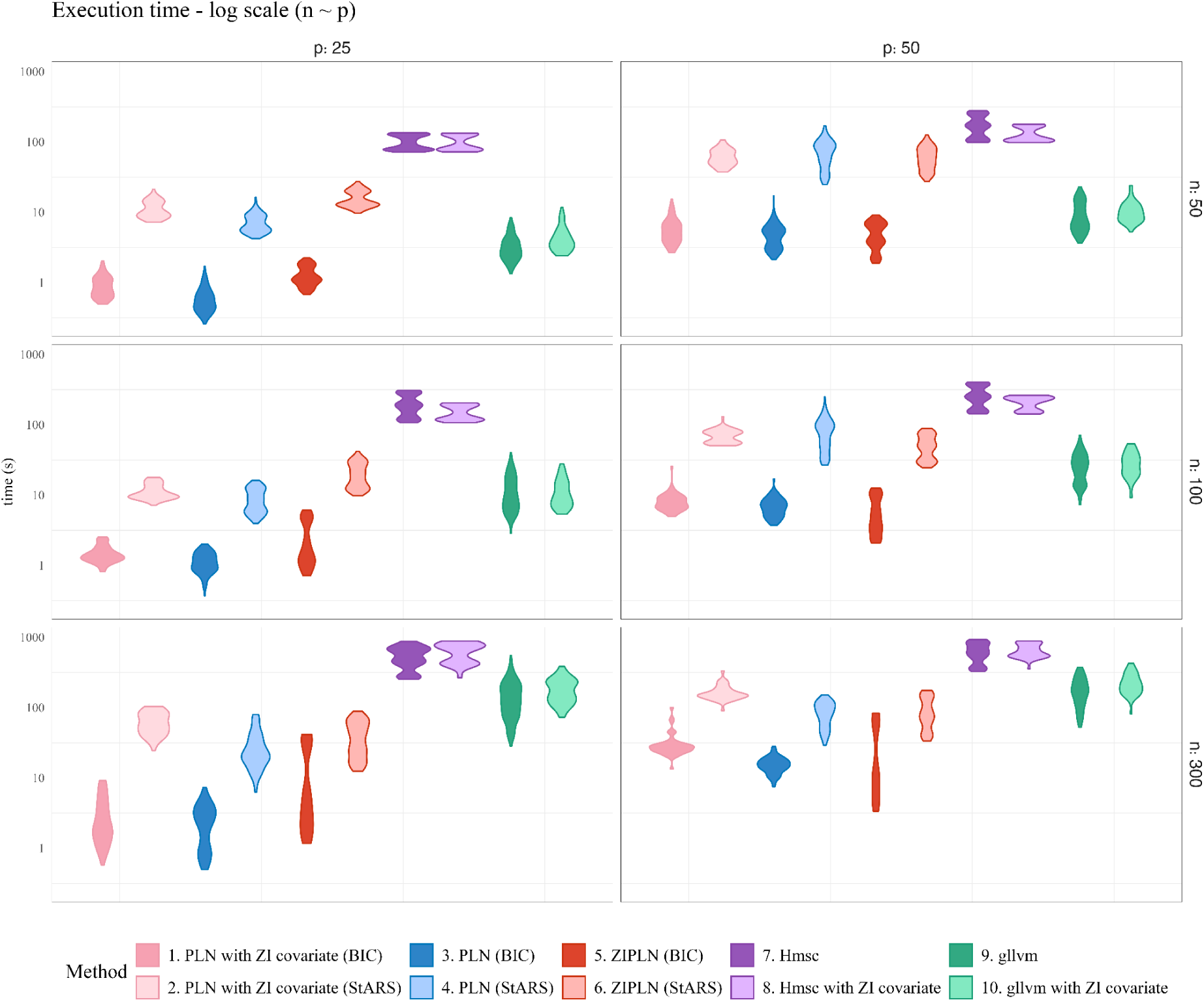
Violin plots of the running times for the different methods and different values of n / p, in logarithmic scale. The zero-inflation is fixed at the reference level (level 3 on the other plots). Using the StARS criterion takes longer because it requires fitting multiple (ZI)PLN models. The Hmsc method requires much more time to run than the others.

### 3.2 Tropical freshwater fish data analysis

A fitting diagnostic for the PLN-network and ZIPLN-network models (Appendix S7) shows that both models fit the data relatively well, with comparable results. Their difference lies in the way they explain the zeros in the data and the consequences it has on the estimation of the partial-correlation network. We compare the association networks (Figure 13) inferred from the tropical freshwater fish data by the PLN-network and ZIPLN-network models with different combinations of covariates (see section 2.6). For a finer interpretation, we include in the analysis the coefficients describing the effects of abiotic covariates on each species that the models infer (Figure 11), corresponding to matrix *B* in the model’s notations (section 2.2) and the zero-inflation probabilities inferred by the ZIPLN-network model (Figure 12). Both give important elements of context to better analyze the species association networks.

**Figure 11:**
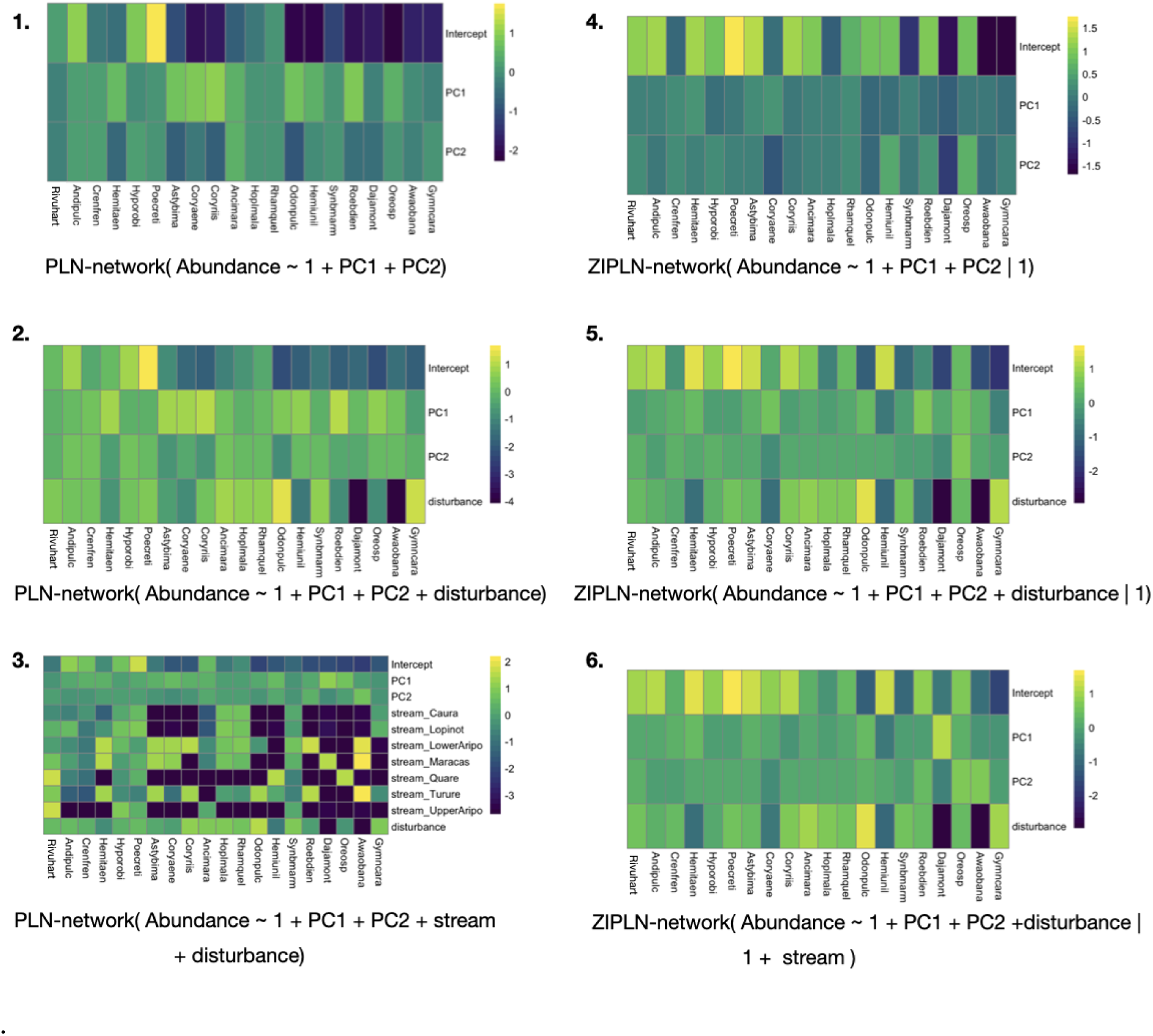
Regression coefficients for abiotic effects on fish abundances across sites (log-scale)

**Figure 12:**
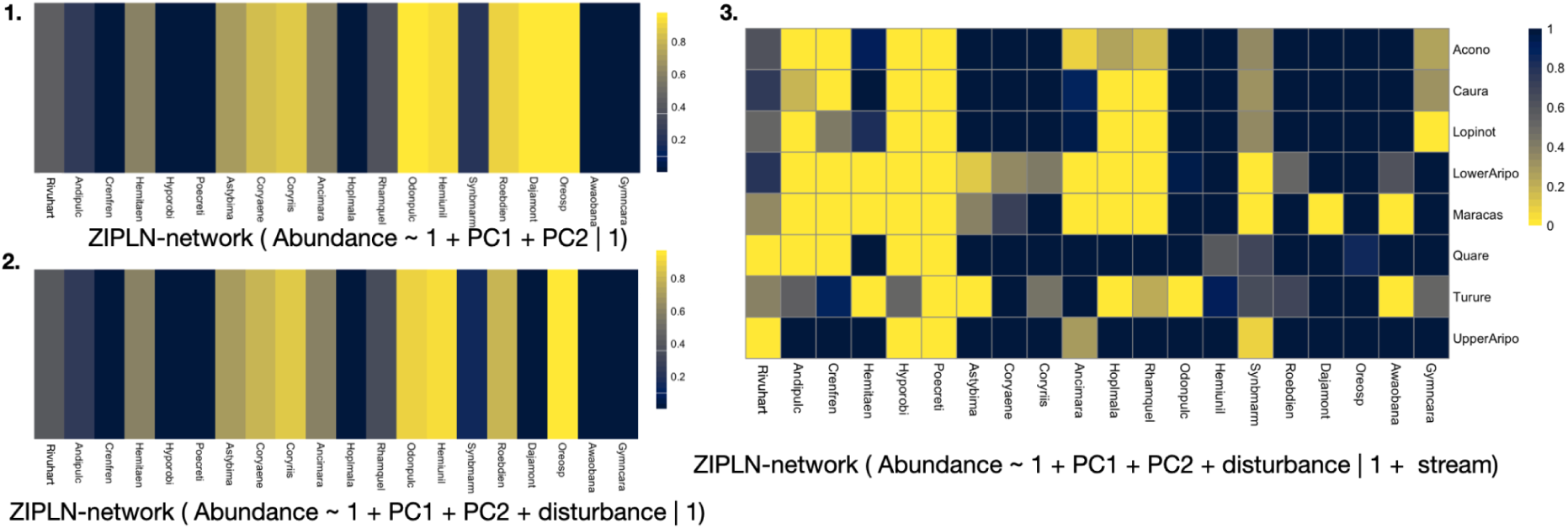
Zero-inflation probabilities inferred by the ZIPLN-network approach with the different combinations of covariates. Plots 1 and 2 correspond to models that consider the zero-inflation probabilities to be species-dependent only whereas this probability depends on both the species and the stream in the third model.

**Figure 13:**
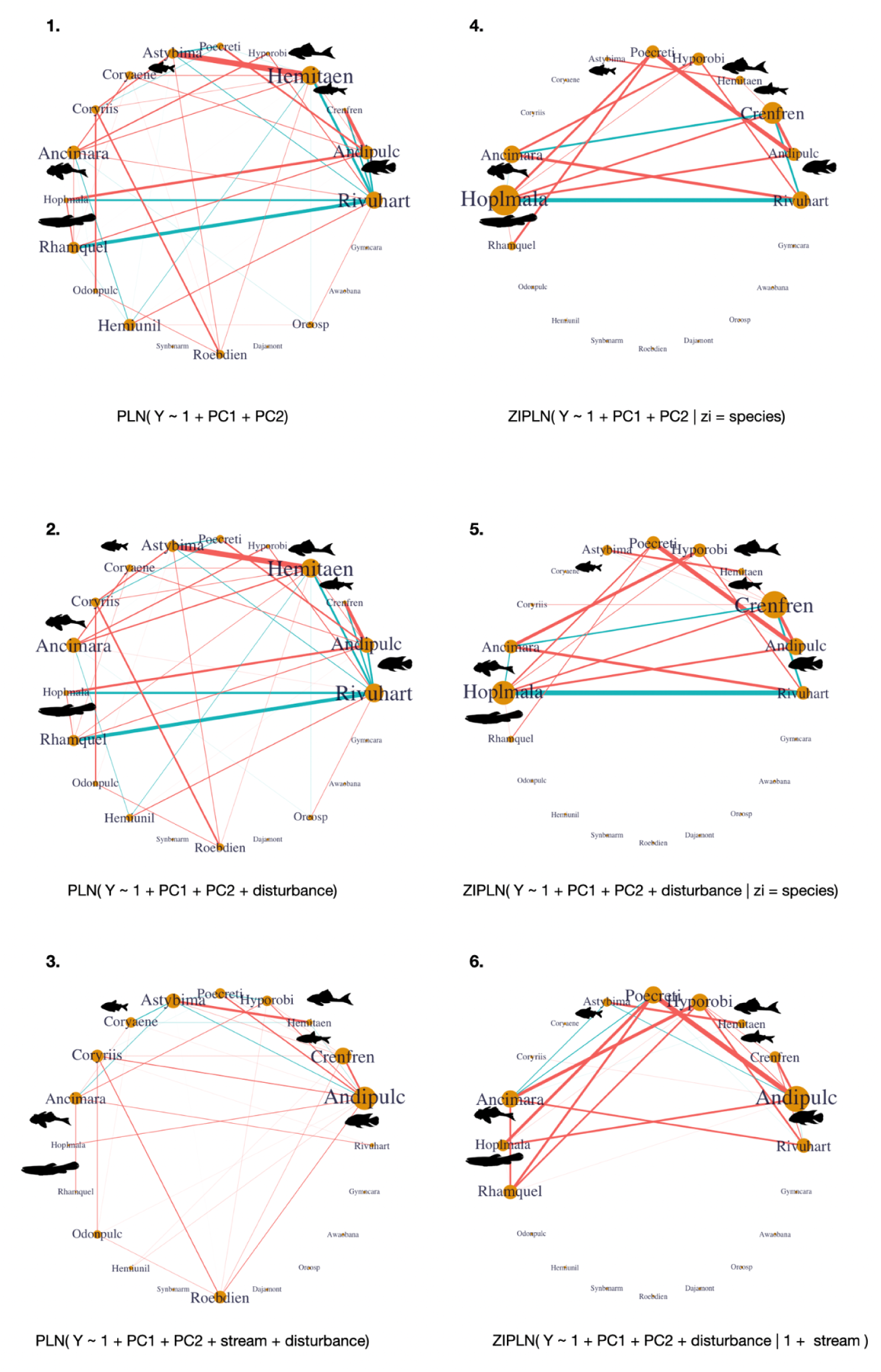
Association networks obtained with PLN-network and ZIPLN-network models for different combinations of covariates. Blue and pink edges represent negative and positive associations, respectively. The thicker the edge, the stronger the association. The size of each species’ name is proportional to the sum of absolute values of associations it is involved in. Silhouettes were retrieved from the Phylopic database. Credits to Grupo de Ictiología de la Universidad de Antioquia. https://creativecommons.org/licenses/by/4.0/

The effects of the environment-related principal components display little variance (Figure 11), with a stronger intercept effect, whereas the stream effect shows an important variance (for the ZIPLN-network model, the stream effect is only included in the zero-inflation coefficients, see Figure 12).

The ZIPLN-network model infers a zero-inflation probability for each pair (*species*, *sample*) depending on the covariates used in the model. Here, for the first two ZIPLN-network models, this probability is only species-dependent whereas for the third one it also depends on the stream considered. When the probabilities are made stream-dependent, for a given species, they can vary a lot from one stream to another (Figure 12).

Accounting for a zero-inflation effect significantly modifies the association networks by removing numerous weak associations (Figure 13). For both the PLN-network and ZIPLN-network models, adding a disturbance effect to abundance only marginally modifies the network. However, accounting for a stream effect, be it in the abundance for the PLN model or in the zero-inflation for the ZIPLN model, removes a certain number of associations. When the stream is accounted for, the PLN and ZIPLN association networks are quite similar, but the associations in the former appear to be much weaker than in the latter case. Some associations are only present in the non-zero-inflated model, such as the positive *Roeboides dientonito* - *Corynopoma riisei* association, the negative associations of *Poecilia reticulata* with both *Astyanax bimaculatus* and *Hypostomus robinii*. Other associations are only visible in the ZIPLN network, such as the positive associations of *Poecilia reticulata* with *Hoplias malabaricus* and *Rhamdia quelen* or the negative one with *Ancistrus maracasae*.

Several associations of interest can also be noted. We find a strong negative association between *Hoplias malabaricus* and *Rivulus hartii* (also known as *Anablepsoides hartii*), present with both approaches, which disappears when the stream variable is taken into account. Positive associations between *Astyanax bimaculatus* and *Hemibrycon taeniurus* on the one hand and *Poecilia reticulata* and *Andinoacara pulcher* on the other hand persist in all the networks.

## 4. Discussion

The new ZIPLN-network approach, introduced in this paper, builds on pre-existing PLN-network and ZIPLN models. We have shown how our approach can be used to analyze zero-inflated, count-based community data in ecology. Our analyses also probed the effects of zero-inflation on the inference of partial-correlation networks in a JSDM context (while taking account of the ways in which different JSDMs compensate for it) and revealed how detrimental such zero-inflation can be when investigating community patterns.

Zeros in species community data can have different origins. Ecologists are often interested in structural zeros resulting from ecological processes, such as the dispersal limits reflected in the freshwater fish species dataset we use as an illustration. We argue that our approach to zero-inflation is particularly useful in these contexts.

### 4.1 Simulations show that the ZIPLN-network model improves the inference of partial-correlation networks in data containing structural zeros

In the simulation study, zeros were added to simulated abundance data, following either species-dependent, site-dependent or covariate-dependent zero-inflation patterns. Such zeros were independent of a species’ rarity when present, thereby emulating structural zeros [Blasco-Moreno et al., 2019].

The AUC results show that failing to model zero-inflation induces a critical degradation of the partial-correlation structure. This outcome persists when considering the inference of marginal correlations. It also appears that the ZIPLN-network model consistently outperforms the reference models in partial-correlation network inference, the focus of this study, but also when considering marginal correlations, although the latter appear harder to retrieve for all the models.

While a penalized model like the ZIPLN-network model can be used to directly obtain a sparse partial-correlation network while applying pre-implemented tools in order to select a given level of sparsity, it appears that integrating penalization in the inference is not critical. Indeed, the ZIPLN model (with no penalty) followed by a penalized inversion of its variance matrix (after the inference procedure) with a Graphical-Lasso, performs just as well as the ZIPLN-network model in terms of AUC (Figures 2 to 4). The advantage of the ZIPLN-network is rather that it directly outputs a sparse precision matrix with no further computing required, and enables the use of specific methods to select the model’s penalty.

Contrasting outcomes between the ZIPLN and the reference models could be explained either by the fact that the ZIPLN model uses a full-rank variance matrix or by other differences in the inference method. A zero-inflated low-rank PLN model is a future step that would extend the comparison presented here, and identify with certainty which hypothesis is more likely.

The simulations also show that increasing the number of observations allows the ZIPLN-network model to largely compensate for increasing zero-inflation, showing that the amount of available data, rather than any bias introduced by zero-inflation, is what limits the ecological insight one can draw from these networks.

Theoretical calculations (Section 2.1) and a detailed example (Figure 5) help understand why modeling structural zeros explicitly through zero-inflation changes the network inferred by the model and can potentially help avoid the detection of spurious associations. Indeed, it shows that shared zero-inflation patterns are interpreted as statistical correlations between species abundances in a non-zero-inflated model whereas they do not appear in the zero-inflated one.

The AUC helps assess the overall performance of each model but it does not resolve the issue of how to best choose the sparsity level of the network through penalty selection in the ZIPLN-network model. The F1-scores (Figure 6) show that the StARS method is better suited than the BIC for penalty selection, although it also requires more time to run (Figure 10). These scores further show that increasing the number of observations is critical.

It is important to note that users of our ZIPLN-network will not be able to extract uncertainty estimates in the current version, as opposed to the reference models. Also, we note that our model does not include random effects either, a useful attribute when the user is interested in the spatial or temporal structure in a JSDM. The Hmsc and gllvm both include these features.

### 4.2 The ZIPLN-network model allows for a refined analysis of the conditional dependencies in the freshwater fish distribution data analysis

In line with what was observed in the simulation study, modeling zero-inflation filters out a number of associations, especially weak ones (Figure 13). In such cases however, one cannot directly tell whether the removed associations were spurious or not. Analysing the association networks considering zero-inflation probabilities can help answer this question. In light of our combined analysis, and given the aforementioned debate regarding the interpretation of statistical correlations as inferred by JSDMs (see Introduction), we stress that our focus is on hypotheses to explain the specific results in our study, remembering that biotic interactions are just one possible explanation for partial correlations [Gotelli et al., 2010].

For instance, the PLN-network model identifies a positive association between *Roeboides dientonito* (hunchback sardine) and *Corynopoma riisei* (swordtail sardine) that is filtered out by the ZIPLN-network model (Figure 13). While this is likely explained by their shared zero-inflation patterns (Figure 6), it is also possibly due to the species sharing the same dispersal limits; the statistical analysis alone cannot rule out the other explanation without the species being present at detectable levels.

Let us now focus on the association network inferred with the ZIPLN-network model using *PC1*, *PC2* (the first two principal components of a PCA run on the covariates that describe each site) and the disturbance level as covariates for abundance, with the stream as a zero-inflation covariate (Figure 13.6).

Several positive associations are found between putative prey and predator species. For instance, *Poecilia reticulata* (guppies) are positively associated with *Andinoacara pulcher* (blue coscorob) and *Hoplias malabaricus* (guabine). Several hypotheses could explain this situation. As guabines are primarily nocturnal predators [Fraser et al., 2006], their coexistence with guppies could reflect temporal niche sharing [Fraser et al., 2004]. Additionally, the behavioural response of guppies to the threat of predation is less in the form of habitat shifts and more likely to be shoaling and avoidance during the most vulnerable crepuscular periods [Fraser et al., 2004]. Guppies and blue coscorobs co-occur in many stream habitats in the Northern Range; the latter is an invertivore, rather than a piscivore, as was sometimes assumed in the earlier literature.

The positive association between *Astyanax bimaculatus* (two-spot sardine) and *Hemibrycon taeniurus* (mountain stream sardine) could be explained by the fact that these species can benefit from adaptive advantages due to heterospecific schooling behaviour [Krause and Ruxton, 2002]. Interestingly, while there is a strong association between these species, there are no other associations within the group of shoaling characids (locally referred to as ‘sardines’) that also includes *Corynopoma riisei* (swordtail sardine) and *Roeboides dientonito* (hunchback sardine). It would be interesting to further explore this pattern to study whether it is explained by different schooling preferences, or by other mechanisms.

While most of the strongest associations seem robust to the addition of new covariates (Figure 13), one striking change is the disappearance of the strong negative association between the apex predator guabine and its prey *Rivulus hartii* (jumping guabine) when a stream covariate is added. This association probably reflects strong predator-avoidance behaviour. Indeed, Fraser and Gilliam (1992) analyzed the predator-prey relationship between both species and found that the presence of guabines suppressed jumping guabine egg production by 50 %. Jumping guabines have excellent jumping abilities (hence their common name) and can survive out of water for extended periods, owing to cutaneous breathing [Fraser, Pers. obs.]. They can easily disperse overland on wet nights to colonise remote bodies of water free of piscivorous fish [Fraser et al. 1995]. These strategies allow them to concentrate at sites that are inaccessible to piscivorous fish. A hypothesis that explains the absence of this association when accounting for a stream effect is the fact that the predator-prey dynamics induce a concentration of jumping guabines in sites where guabines cannot be found, a pattern that is caught in the zero-inflation, stream-related part of the model. Looking back at the abundance heatmap (Figure 1), it indeed appears that jumping guabines seem to thrive more at sites where guabines are completely absent.

This example raises the question of identifiability in the effects inferred by the different models as, if our hypothesis correctly explains the presence and disappearance of this association, then it means that a predator-prey avoidance strategy is dismissed from the network inferred with zero-inflation. We argue that it is precisely the combined analysis of the zero-inflated and non-zero-inflated model that allows one to formulate a finer hypothesis regarding this phenomenon that is probably explained by both biotic interactions and the presence of dispersal barriers.

Thus, as we have shown how, in this example, the ZIPLN-network analysis allows one to propose several hypotheses regarding the biotic and abiotic mechanisms that shape community structure. It can also be used to understand how pre-identified biotic interactions show up in community data. Moreover, comparing the ZIPLN-inferred network with its non-zero-inflated counterpart helps formulate better-informed hypotheses regarding the ecological processes at play.

## 5. Conclusion

We conclude that the ZIPLN-network model is a useful tool for the analysis of correlation networks, and specifically partial-correlation networks, in zero-inflated community data. Our simulation study shows that accounting for the presence of structural zeros (zeros that emanate from the zero-inflation in a simulation context) is critical to correctly infer partial-correlation networks from zero-inflated count data (with the correct associations being known in a simulation context) and that the ZIPLN-network model offers a useful way to do so, provided it is given enough observations.

We have also shown that comparing the results obtained with the PLN-network and ZIPLN-network models helps better disentangle the assemblage mechanisms at stake. Finally, it can help distinguish dispersal effects from other mechanisms by accounting for the absence of species at sites they did not reach.

In conclusion, we have demonstrated how the ZIPLN-network approach can be used to produce more informed analyses of community data. Accounting for zero-inflation and using both presence-absence and count data can filter out spurious associations and help distinguish different mechanisms. We argue that such a model is ideally used in combination with its non-zero-inflated counterpart so as to better identify putative ecological mechanisms. Similarly, comparing the results obtained with different combinations of covariates allows for a more nuanced analysis.

## Acknowledgements

This work was supported by public grants from the Fondation Mathématique Jacques Hadamard, AgroParisTech, the DESSE INRAE and the Erasmus+ program. We thank Stéphane Robin and Wilfried Thuiller for the fruitful discussions about this work.

## Author contributions

JT led the investigation. JT, JC & AEM designed the study. JC designed the ZIPLN-network model and provided technical expertise on its use. JT ran the analyses. AEM, AED, A F-E & DFF provided ecological expertise to support the interpretation of the results. JT wrote the initial draft. All authors reviewed and approved the final manuscript.

License details for species silhouettes that are not in the public domain:

For Ancistrus maracasae, Andinoacara pulcher, Hemibrycon taeniurus, Hoplias malabaricus, Hypostomus robinii, Roeboides dientonito,: credits to Grupo de Ictiología de la Universidad de Antioquia. https://creativecommons.org/licenses/by/4.0/

For Rhamdia quelen, Crenicichla frenata: credits to Cesar Julian. https://creativecommons.org/licenses/by/3.0

## Appendices

## Appendix S1: Simulation study under a sparse variance-covariance matrix

In this appendix are presented the results of a simulation study run under the same conditions as the main simulation study of the manuscript, with the difference that it is the variance matrix that was simulated as sparse instead of the precision matrix. We study how the models perform at retrieving the support of this matrix.

**Figure S1.1:**
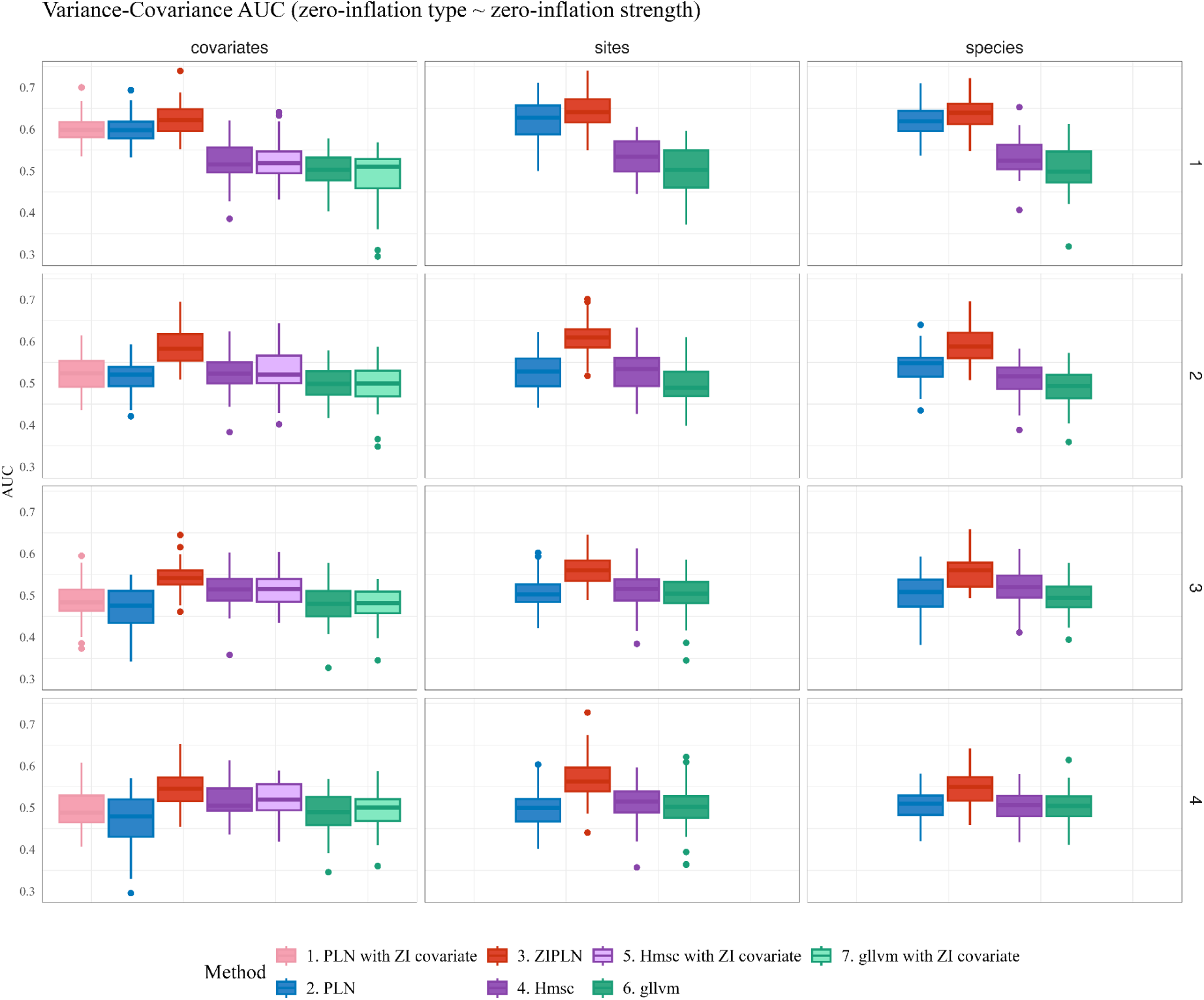
Boxplot of the AUCs obtained for the variance matrix with the different network inference methods tested, as the zero-inflation type and zero-inflation level (1 = lowest, 4 = highest) vary, with

**Figure S1.2:**
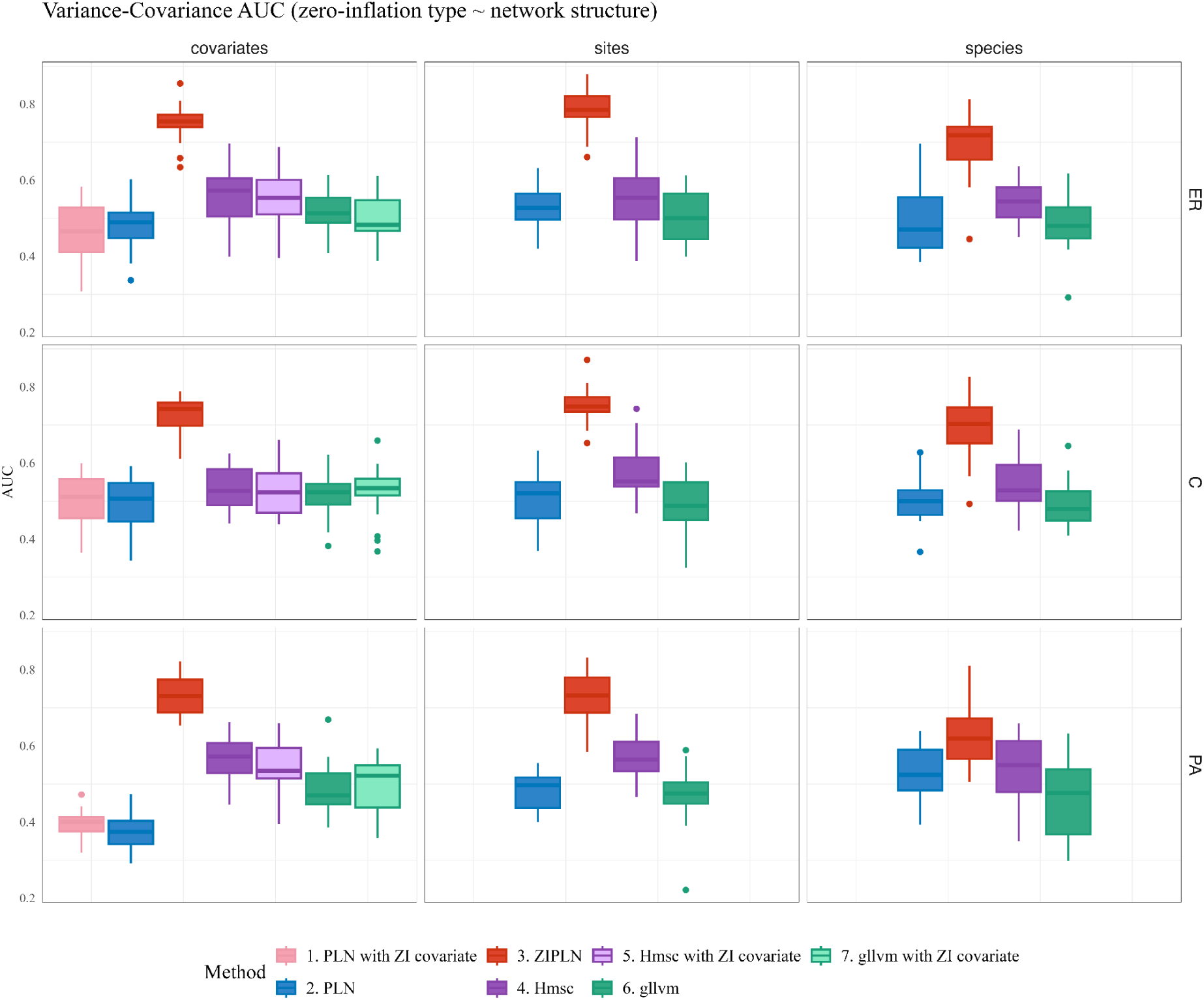
Boxplot of the AUCs obtained for the variance matrix with the different network inference methods tested as the network structure and zero-inflation type vary for a zero-inflation fixed at the reference level (level 3 on the other plots), with *n* = 100 and *p* = 50 (ER: Erdös-Rényi, PA: Preferential Attachment, C: Community).

**Figure S1.3:**
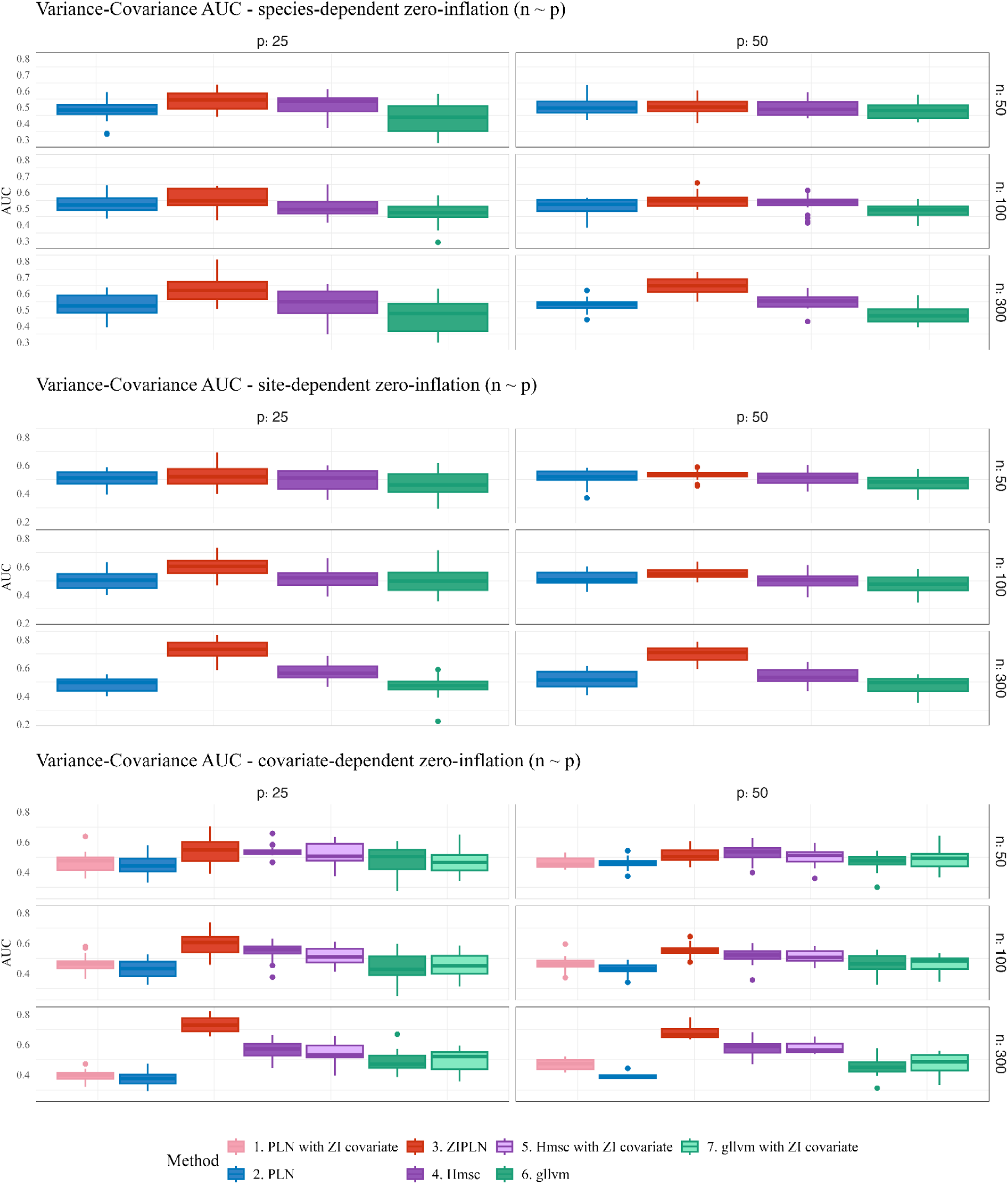
Boxplot of the AUCs obtained for the variance matrix with the different network inference methods tested as the number of sites (*n*) and of species (*p*) vary for the different zero-inflation types, with a zero-inflation level fixed at the reference level (level 3 on the other plots).

**Figure S1.4:**
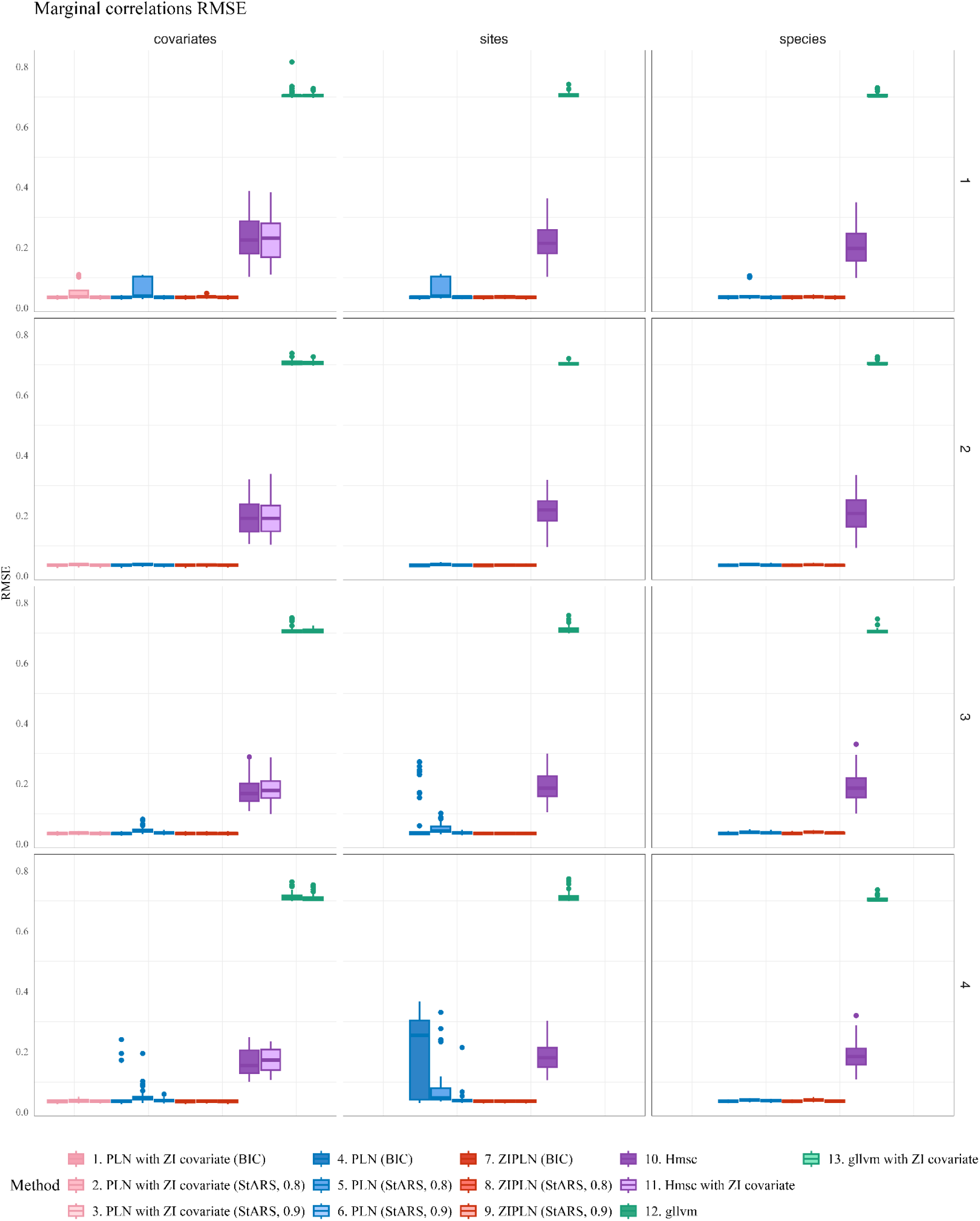
Boxplot of the RMSE obtained for the marginal correlations by each method, with each criterion when applicable, for each zero-inflation configuration (zero-inflation levels: 1 = lowest, 4 = highest), with *n* = 100 and *p* = 50. Each boxplot corresponds to 60 replicates (20 for each of the three tested network structures).

**Figure S1.5:**
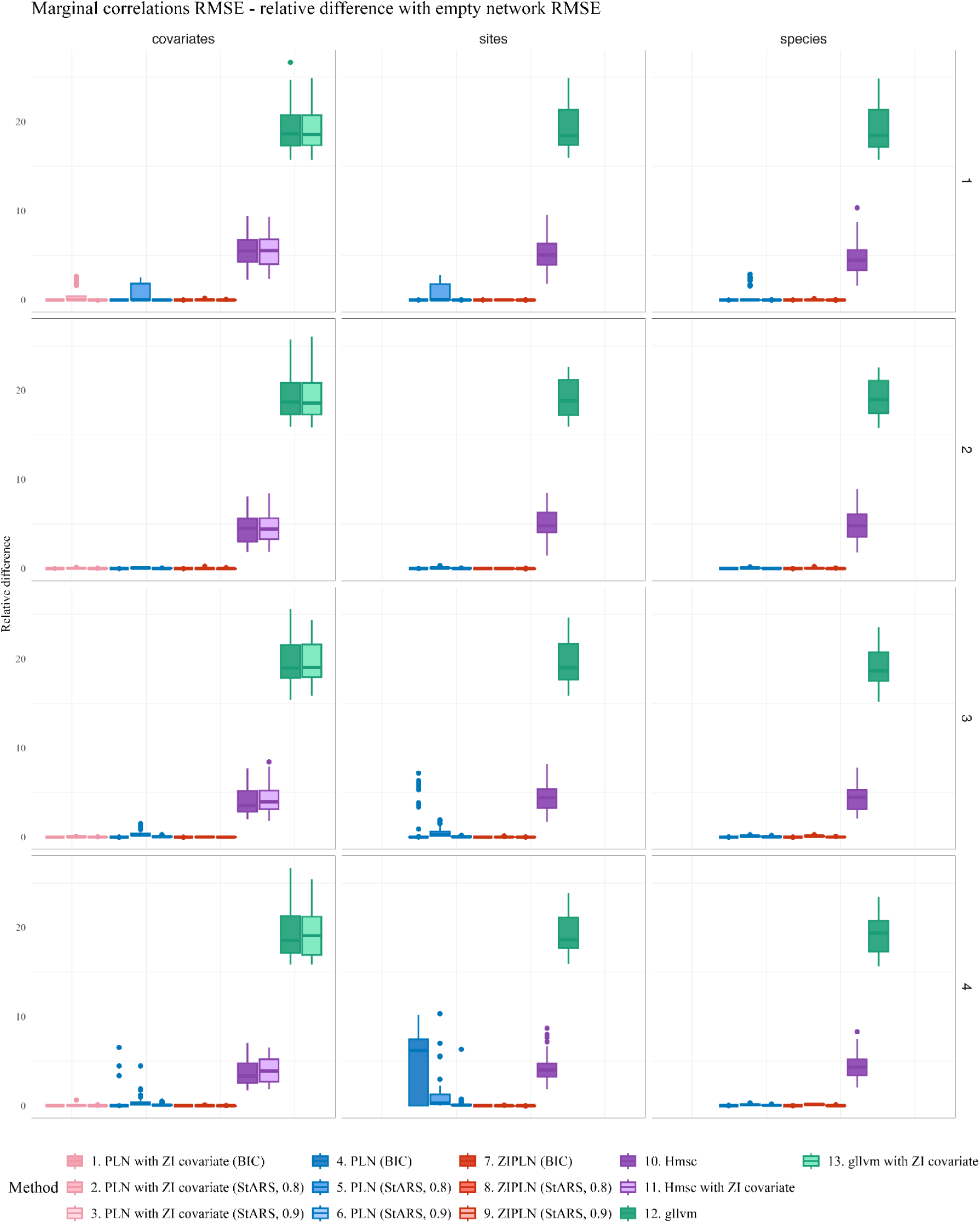
Relative difference between the marginal correlations RMSE obtained by each method and each criterion (if applicable) and the RMSE obtained with an empty network (all partial correlations equal to 0), with *n* = 100, *p* = 50 (zero-inflation level: 1 = lowest, 4 = highest). Each boxplot corresponds to 60 replicates (20 for each of the three tested network structures).

## Appendix S2: Testing the gllvm and Hmsc model with increased ranks

We tested whether the gllvm and Hmsc results could be improved by increasing the number of latent factors. For gllvm the manuscript’s simulations were run with 2 latent factors whereas the Hmsc model determined how many latent factors should be used. For this supplementary material, we ran both models with a number of latent factors going from 2 to 7. We measured the results over 20 replicates of a simulation under the standard protocol we have used throughout the study.

The AUC, running times and situations of failed convergence were measured for both models. Regarding convergence, the gllvm model directly provides an indication of convergence. For the Hmsc model, we ran the optimization with two chains and checked the convergence with the potential scale reduction factor across the chains (Gelman and Rubin, 1992). We considered that the model had converged if the median of the potential scale reduction factor was below 1.1 for both the regression and variance parameters.

For the gllvm model, increasing the number of latent variables did not improve network recovery in practice: beyond four latent variables the gllvm optimizer failed to converge at these dimensions (*n* = 100, *p* = 25), while the fitting time increased by two orders of magnitude. We report this as a practical limitation of the fitting procedure rather than of the low-rank representation itself. The Hmsc model systematically converged, and while the running time increased, this remained reasonable. The AUCs, however, did not improve either with the number of latent factors.

**Figure S2.1:**
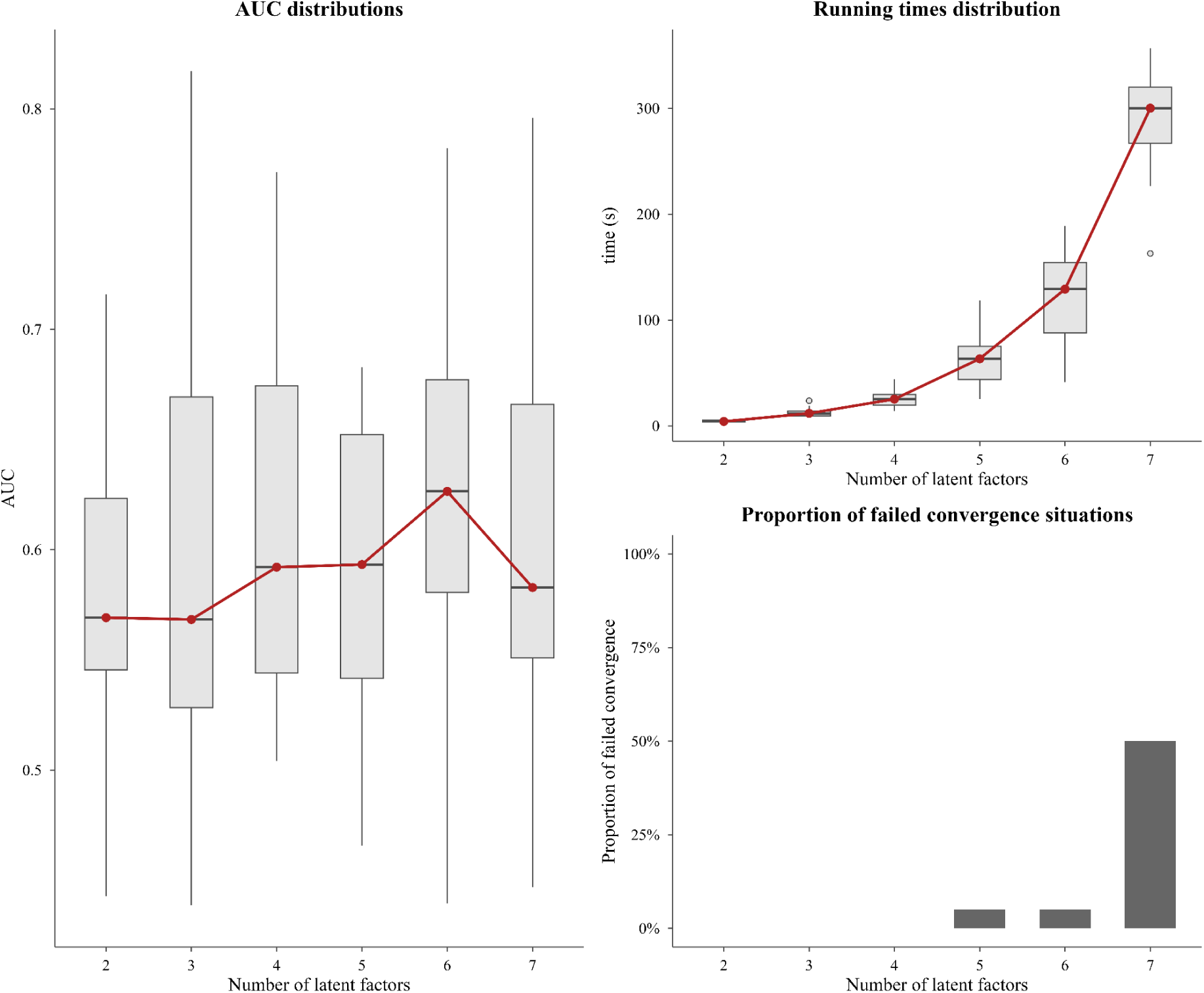
gllvm results when increasing its number of latent factors, over 20 replicates of a simulation with *n* = 100, *p* = 25, an Erdös-Rényi network structure and a species-dependent zero-inflation at the reference level. **Left:** distribution of the partial-correlation network AUCs. **Right - top:** running times. **Right - bottom:** proportion of situations in which the model convergence failed.

**Figure S2.2:**
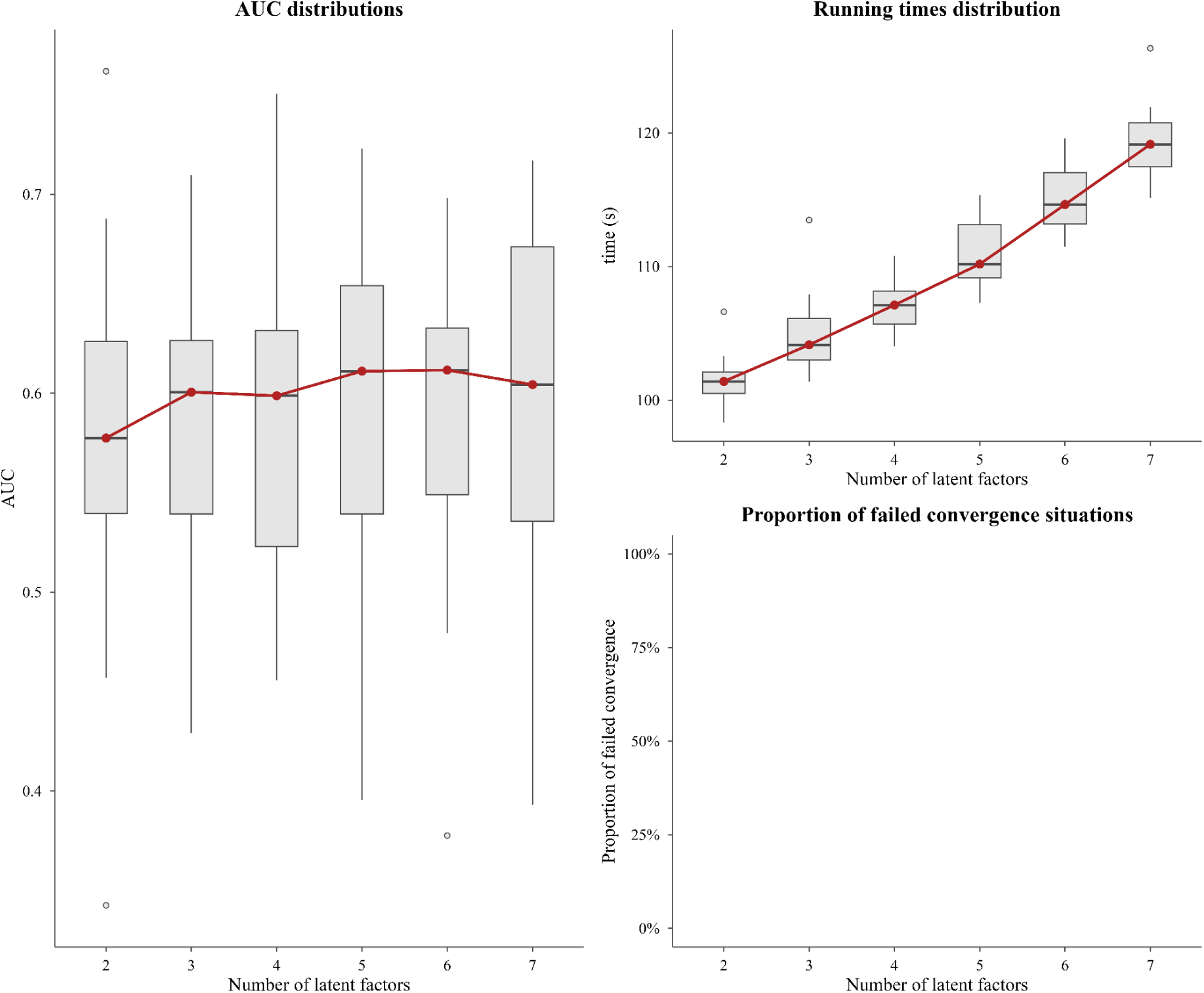
Hmsc results when increasing its number of latent factors, over 20 replicates of a simulation with *n* = 100, *p* = 25, an Erdös-Rényi network structure and a species-dependent zero-inflation at the reference level. **Left:** distribution of the partial-correlation network AUCs. **Right - top:** running times. **Right - bottom:** proportion of situations in which the model convergence failed (according to the Gelman and Rubin criterion we used, no convergence issue arose).

## Appendix S3: Simulation study under a negative-binomial distribution

**Figure S3.1:**
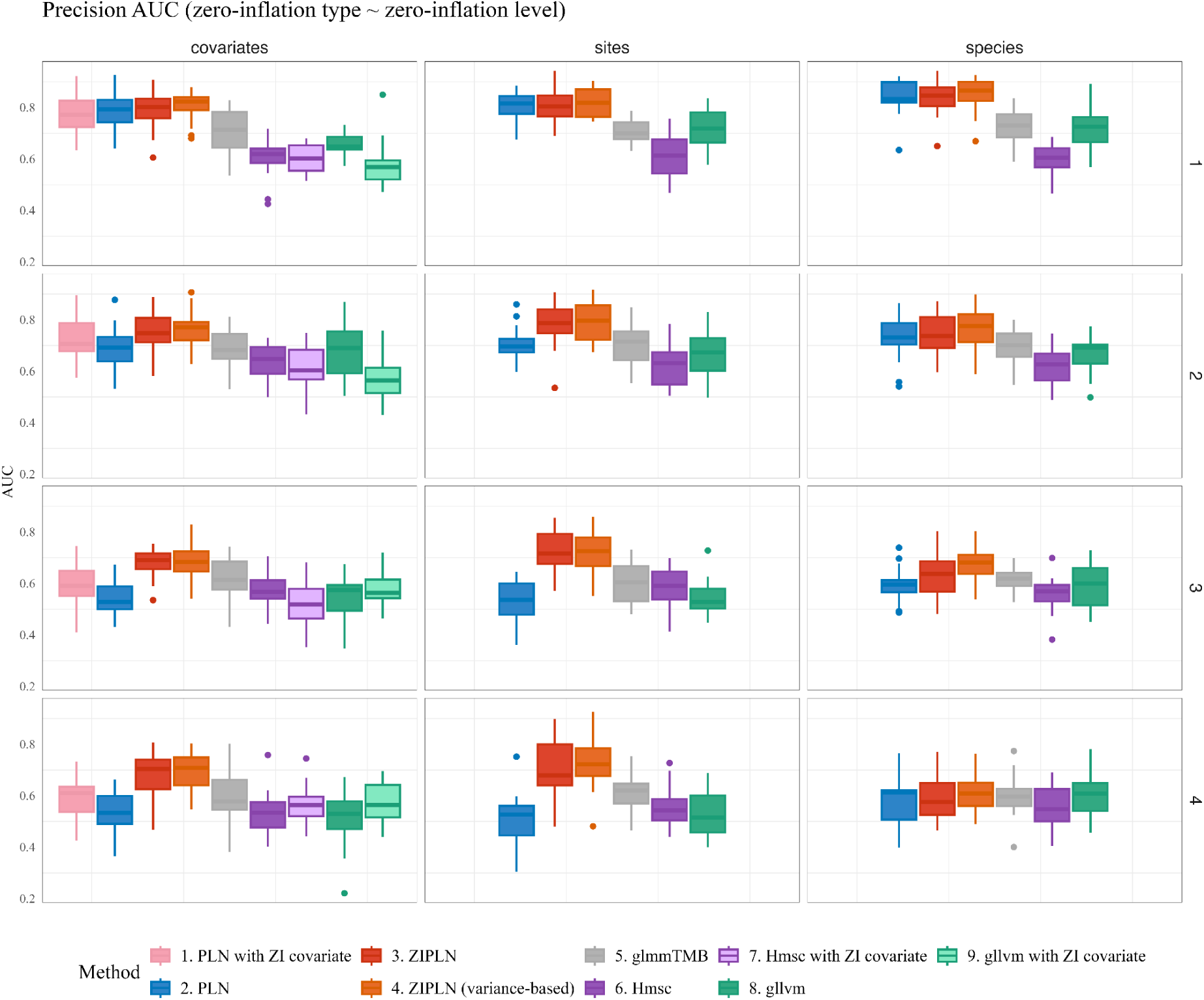
Boxplot of the AUCs obtained with the different network inference methods tested, as the zero-inflation type and level vary when data is simulated under a negative-binomial distribution, with *p* = 25, *n* = 100. Each simulation has 20 replicates. The zero-inflation levels are denoted 1 to 4 from lowest (few structural zeros) to highest (many structural zeros).

**Figure S3.2:**
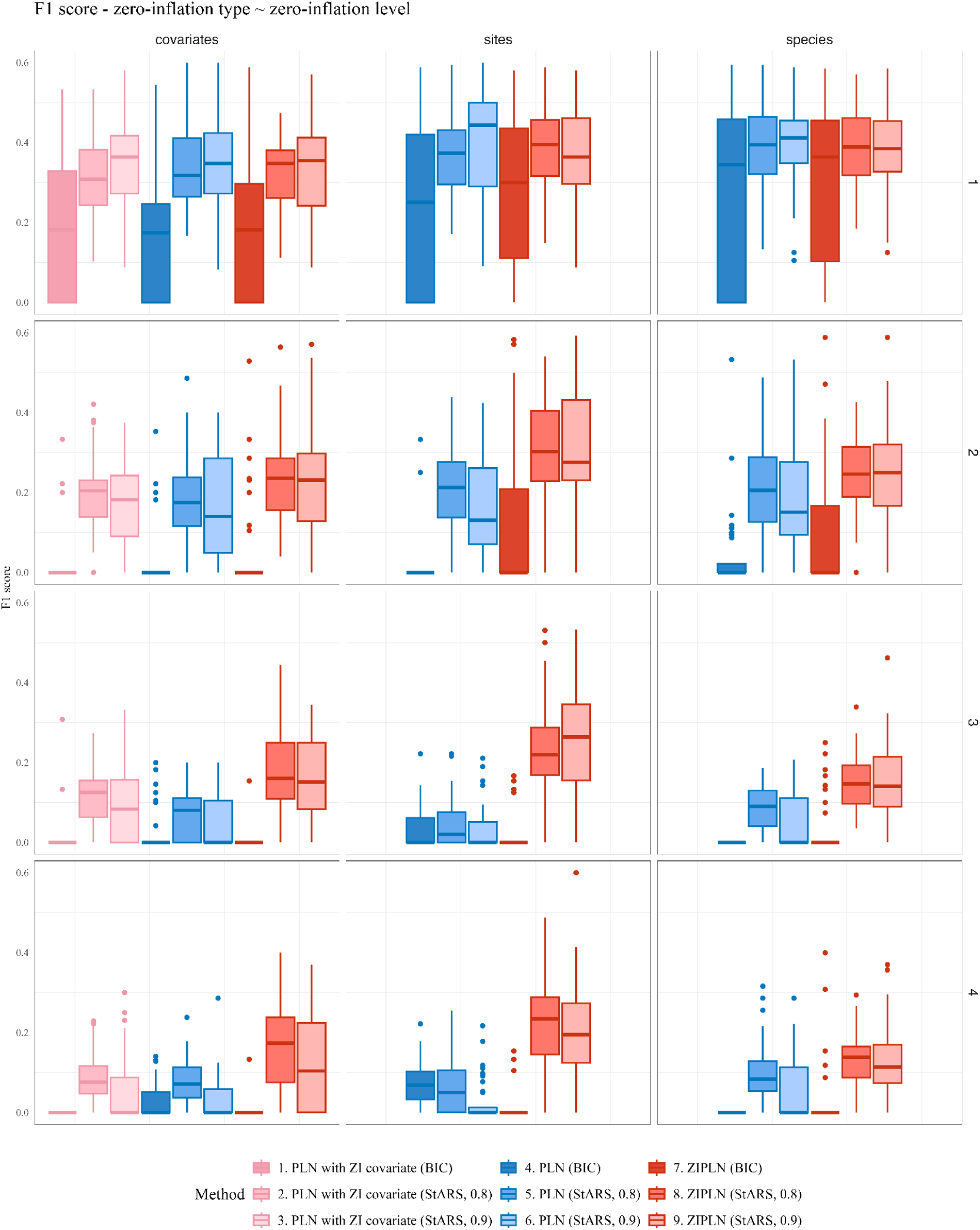
Boxplot of the F1 scores obtained with the PLN and ZIPLN methods for penalties selected with different criteria (either BIC or StARS at levels 0.8 or 0.9), for different zero-inflation types and levels, when data is simulated under a negative-binomial distribution, with *p* = 25, *n* = 100. Each simulation has 20 replicates. The zero-inflation levels are denoted 1 to 4 from lowest (few structural zeros) to highest (many structural zeros).

**Figure S3.3:**
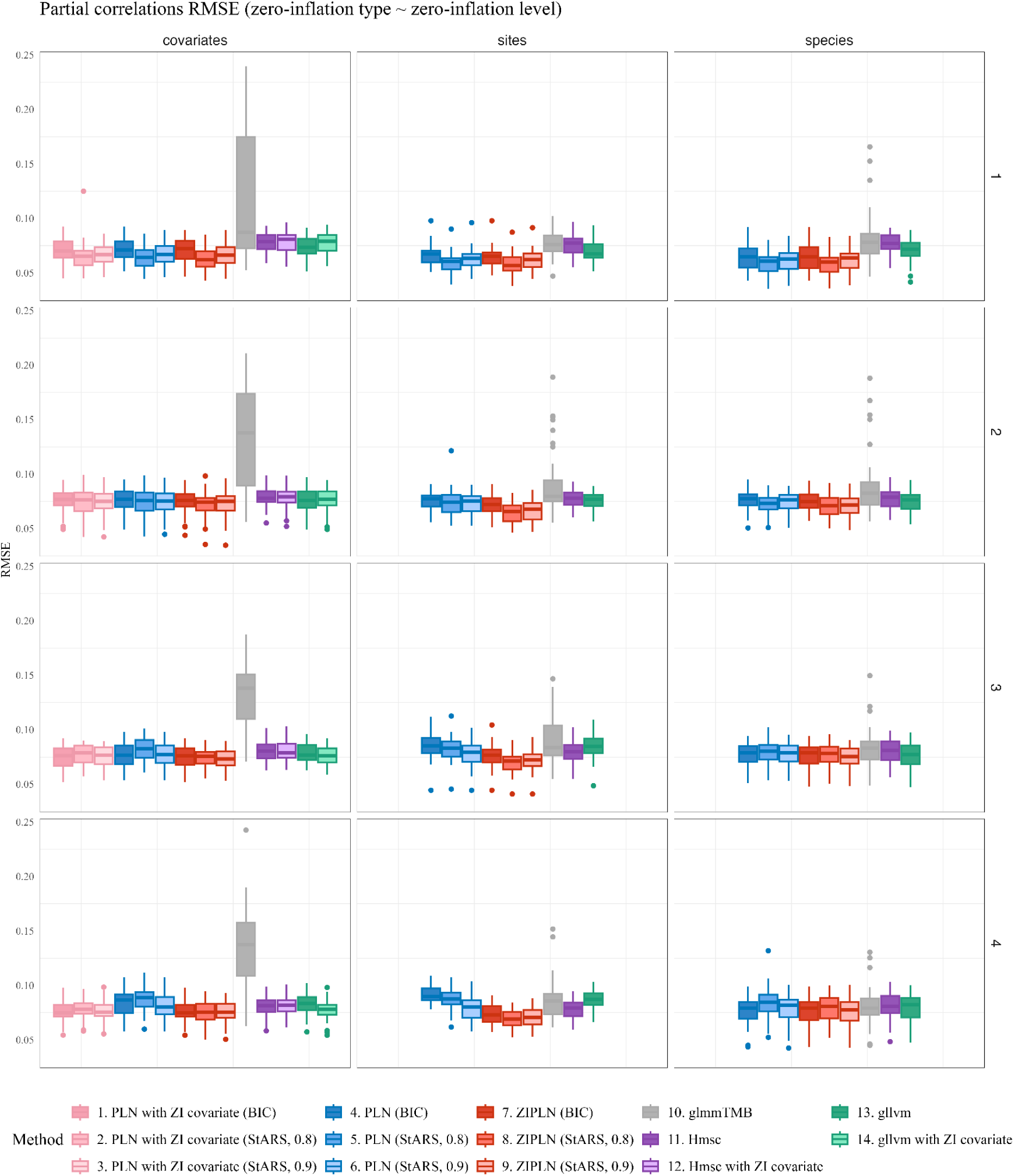
Boxplot of the RMSE obtained for the partial correlations by each method, with each criterion when applicable, for each zero-inflation configuration,when data is simulated under a negative-binomial distribution, with *p* = 25, *n* = 100. Each simulation has 20 replicates. The zero-inflation levels are denoted 1 to 4 from lowest (few structural zeros) to highest (many structural zeros).

**Figure S3.4:**
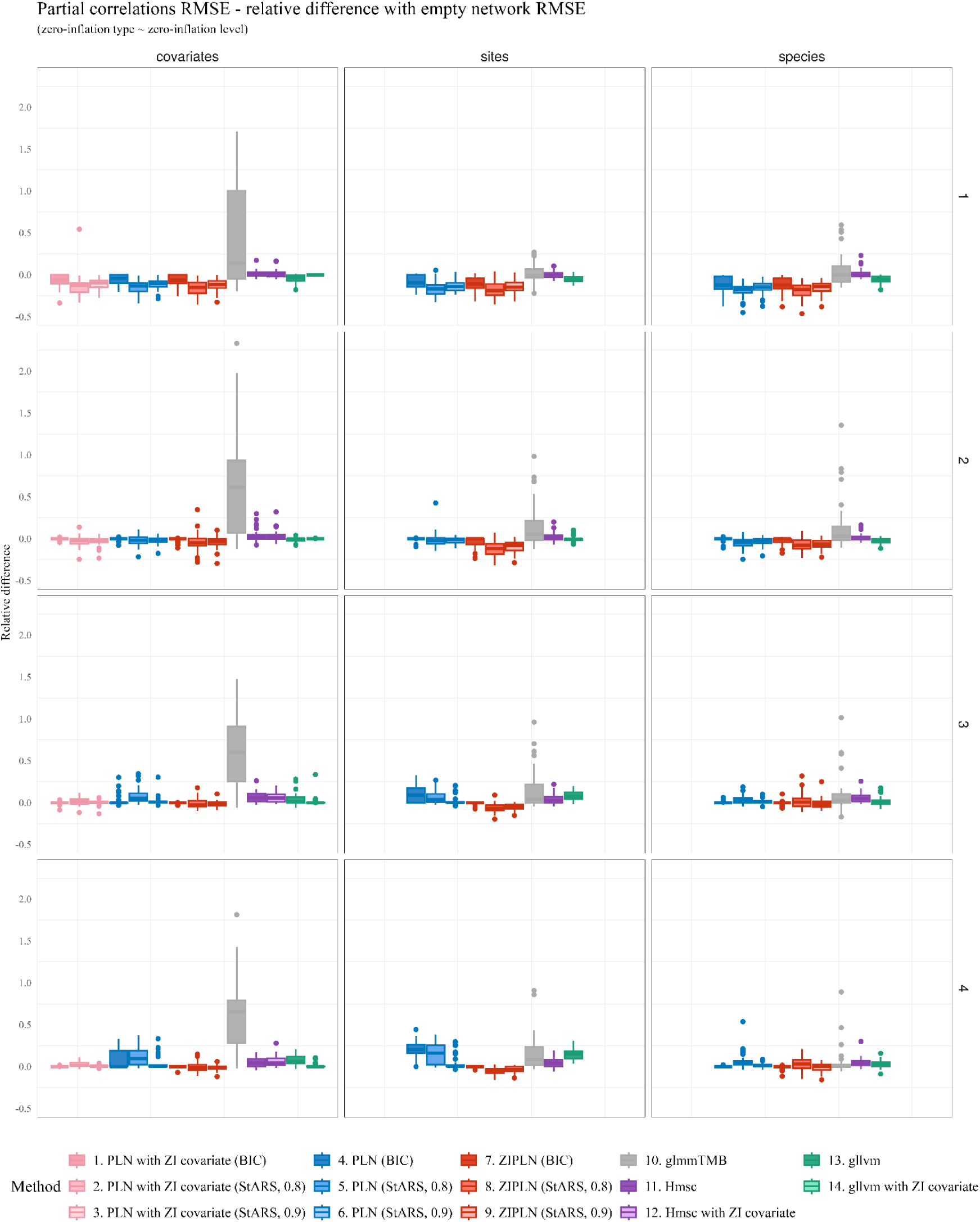
Relative difference between the partial correlations RMSE obtained by each method and each criterion (if applicable) and the RMSE obtained with an empty network (all partial correlations equal to 0), when data is simulated under a negative-binomial distribution, with *p* = 25, *n* = 100. Each simulation has 20 replicates. The zero-inflation levels are denoted 1 to 4 from lowest (few structural zeros) to highest (many structural zeros).

**Figure S3.5:**
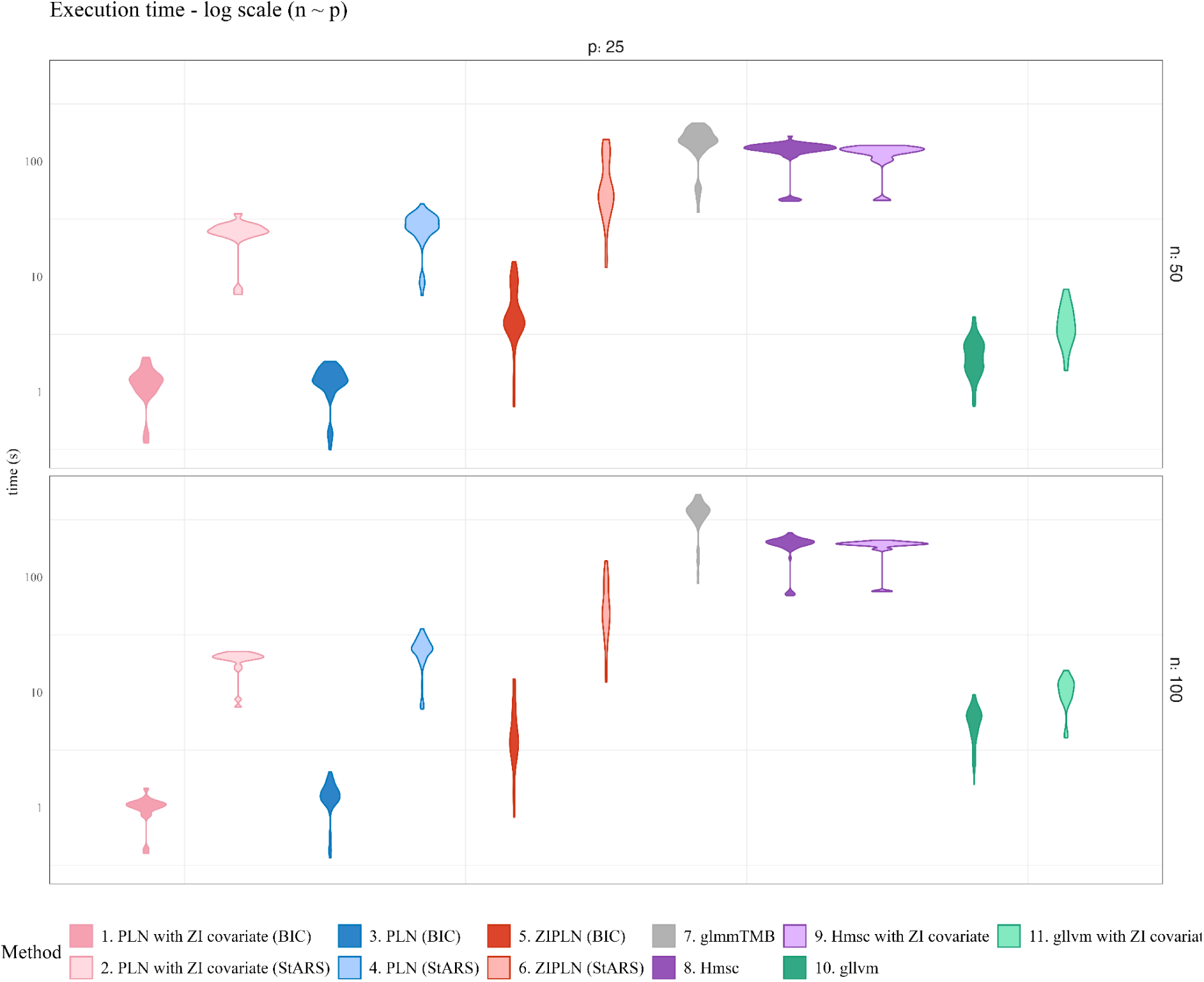
Execution times of the different methods when data is simulated under a negative-binomial distribution and the zero-inflation is set at the reference level (level 3 on the other Figures) for different values of *n* and *p*. Each simulation has 20 replicates.

## Appendix S4: Correspondence between full species names and their shortened versions

**Table S4:**
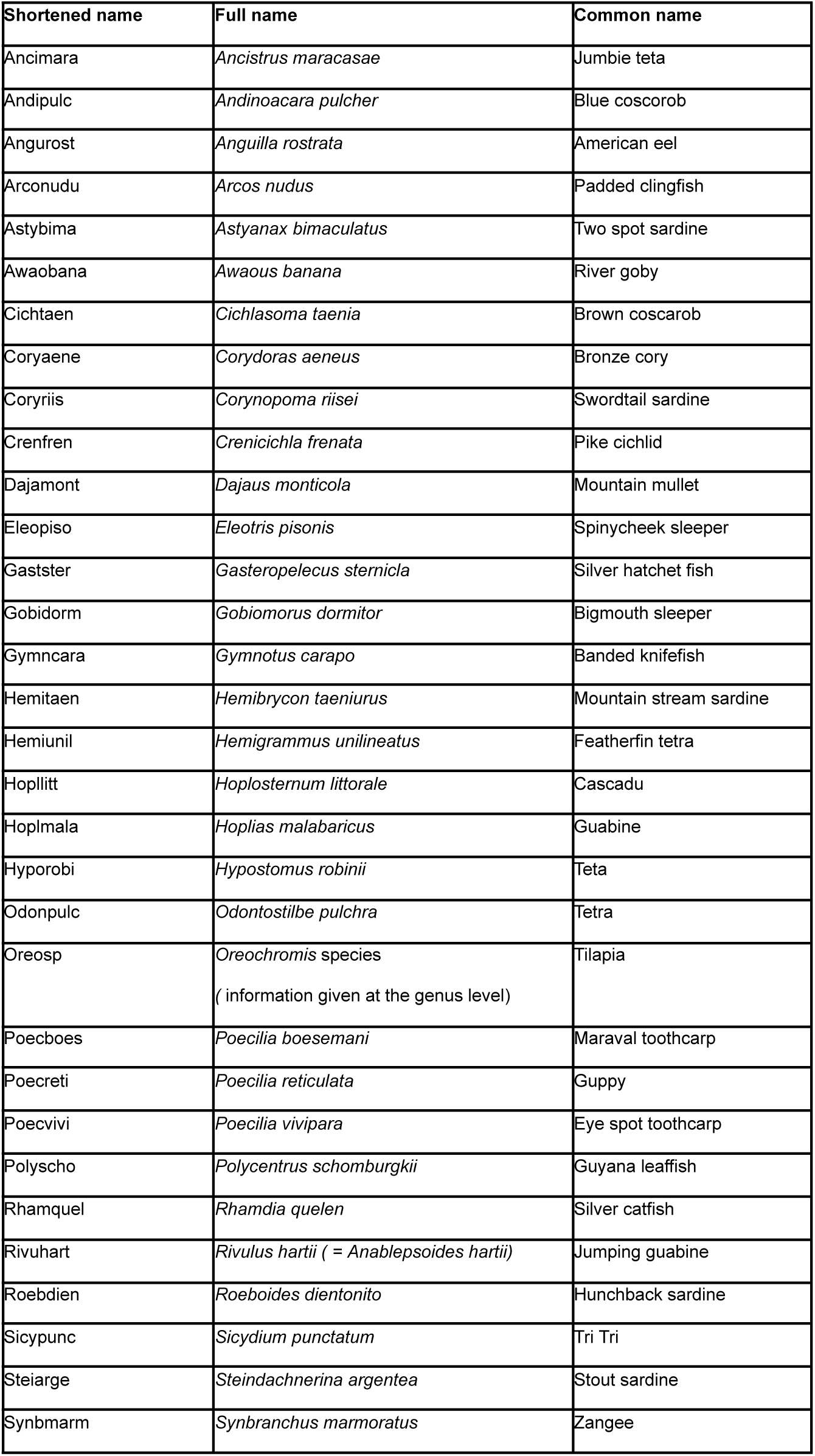
Correspondence between full species names and their shortened versions.

## Appendix S5: Results of the Principal Component Analysis on the environmental covariates

**Figure S4:**
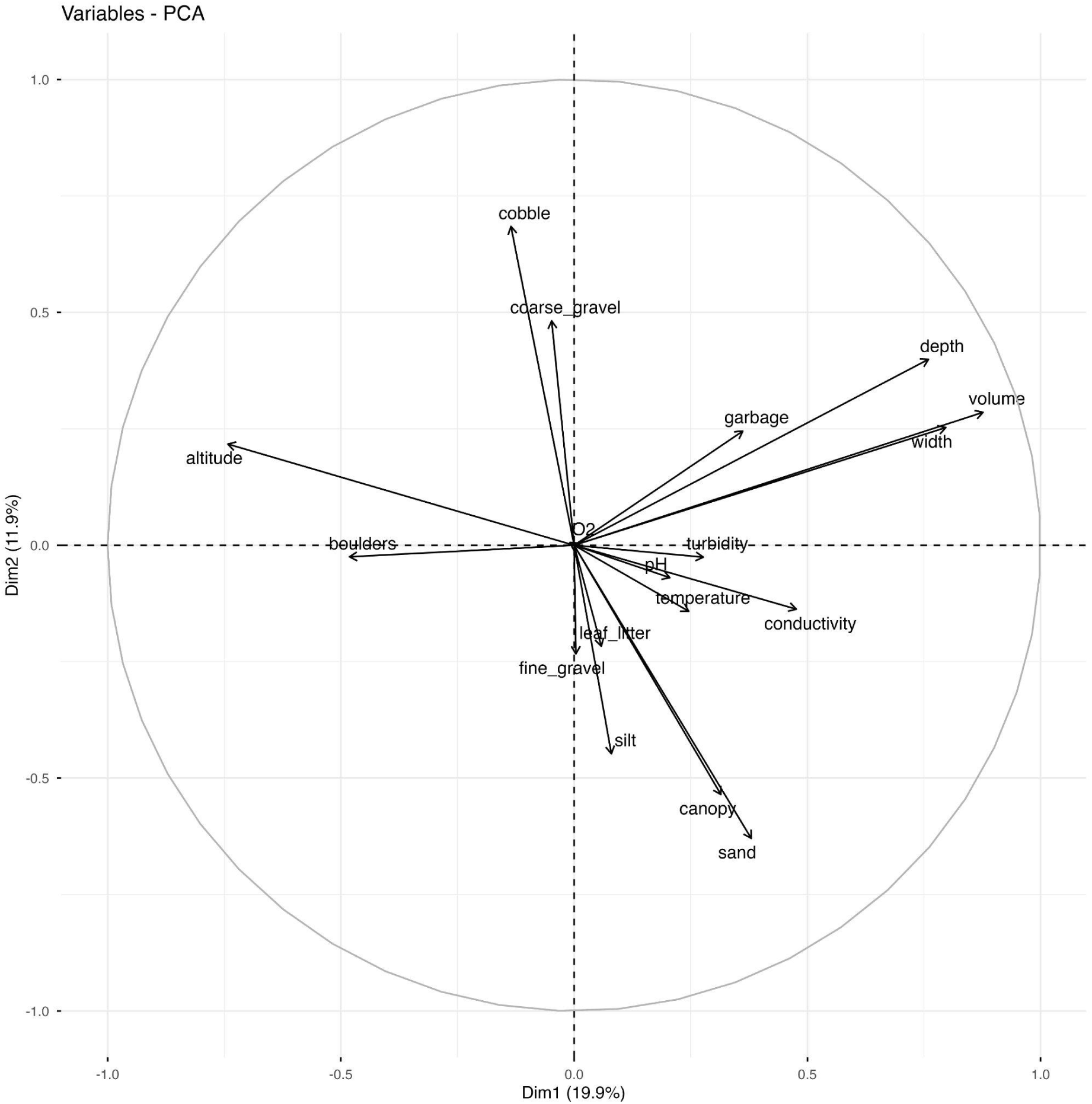
Results of the Principal Component Analysis on the environmental covariates.

This figure describes the weight of each covariate in each of the first two principal components resulting from the PCA run on the tropical freshwater fish species distribution dataset. The variables used include each site’s altitude, the depth, width, volume, temperature, turbidity, conductivity, oxygen concentration and pH of the water, descriptors of the canopy and of the soil substrates (boulder, fine gravel, leaf litter, sand, leaf litter…).

## Appendix S6: A quick guide to using the ZIPLN-network model for the analysis of species abundance data containing structural zeros

The code used for the study of the analysis of the tropical freshwater fish distribution data is provided in the github repository: jeannetous/ZI_JSDM_for_association_network_inference_from_ZI_count_based_community_data

For more details about the R package **PLNmodels** and its features, the reader may refer to [Chiquet et al., 2021] or to https://pln-team.github.io/PLNmodels/articles/.

We provide a quick step-by-step guide to analyze species distribution data with structural zeros using the R package **PLNmodels**. This guide does not include all the refinements detailed in the **PLNmodels** documentation. The user should first install and load the **PLNmodels** package (version 1.2.2 or higher).

1. **Data formatting:**

- Species distribution data must be turned into a matrix *Y* with *n* rows (the number of sites) and *p* columns (the number of species), with *Y_ij_* an integer giving the number of individuals from species *j* in sites *i*. If the data are contained in a csv, this can be done using the **read.csv** R function.
- The covariates must be turned into a matrix *X* with *n* rows (the number of sites) and *d* columns (the number of covariates). These covariates can be the results of a PCA or other transformations of the original covariates; we used a PCA (section 2.6 and Appendix S4).
- A data object **df** specifically designed for the **PLNmodels** functions is then created using the function **prepare_data** of **PLNmodels**: **df** ← **prepare_data(Y, X)**.
2. **Running the ZIPLN-network model:**

- The user must first decide which covariates should be included in the model. This can be done *a priori* using ecological knowledge of the data or *a posteriori* comparing statistical criteria describing the model fitting quality obtained with different covariates combinations. Then denote *X*_1_ to *X_d_* the selected covariates. We consider that an intercept is also used. In our study we selected as covariates the first two components of a PCA run on a number of environmental covariates (see section 2.6).
- The user should also decide what type of zero-inflation is required: sites, species or covariates-dependent and, in the latter case, decide what covariates should be used. We denote them 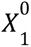 to 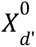. In our study, after analysing the data (Figure 1) we tested both a species-dependent zero-inflation and a covariate-dependent zero-inflation, using the stream covariate (see section 2.6).
- For a species-dependent zero-inflation, the user should run:

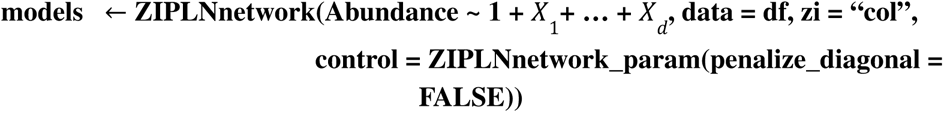
- For a covariates-dependent zero-inflation (including an intercept), the user should run:

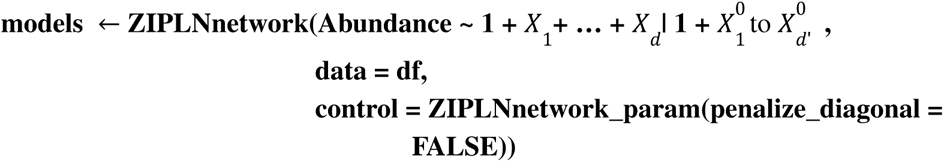
3. **3. Analyzing the results:**

- The object **models** contains several ZIPLN-network models, one per *P* penalty used (see section 2.2 and [Chiquet et al., 2019]). The user must first select one model, either picking a specific penalty λ running **model** ← **models$getModel (** λ) or by selecting the penalty that gives the best result for a specific criterion (BIC, EBIC or StARS) **model** ← **models$getBestModel(*criterion***). In our study, we used the StARS selection criterion with a stability selection value of 0.8 (see section 2.2).
- The user may then access several attributes of the object **model** to study the results. Details are given in the **PLNmodels** documentation but the user may for instance visualize the inferred species association network running **model$plot_network()** (Figure 13) or extract the regression coefficients using **model$model_par$B** (Figure 11) or the zero-inflation probabilities with **model$model_par$Pi** (Figure 12).

## Appendix S7: Fitting diagnostics of the PLN and ZIPLN model for the Trinidad fish community dataset

**Figure S7:**
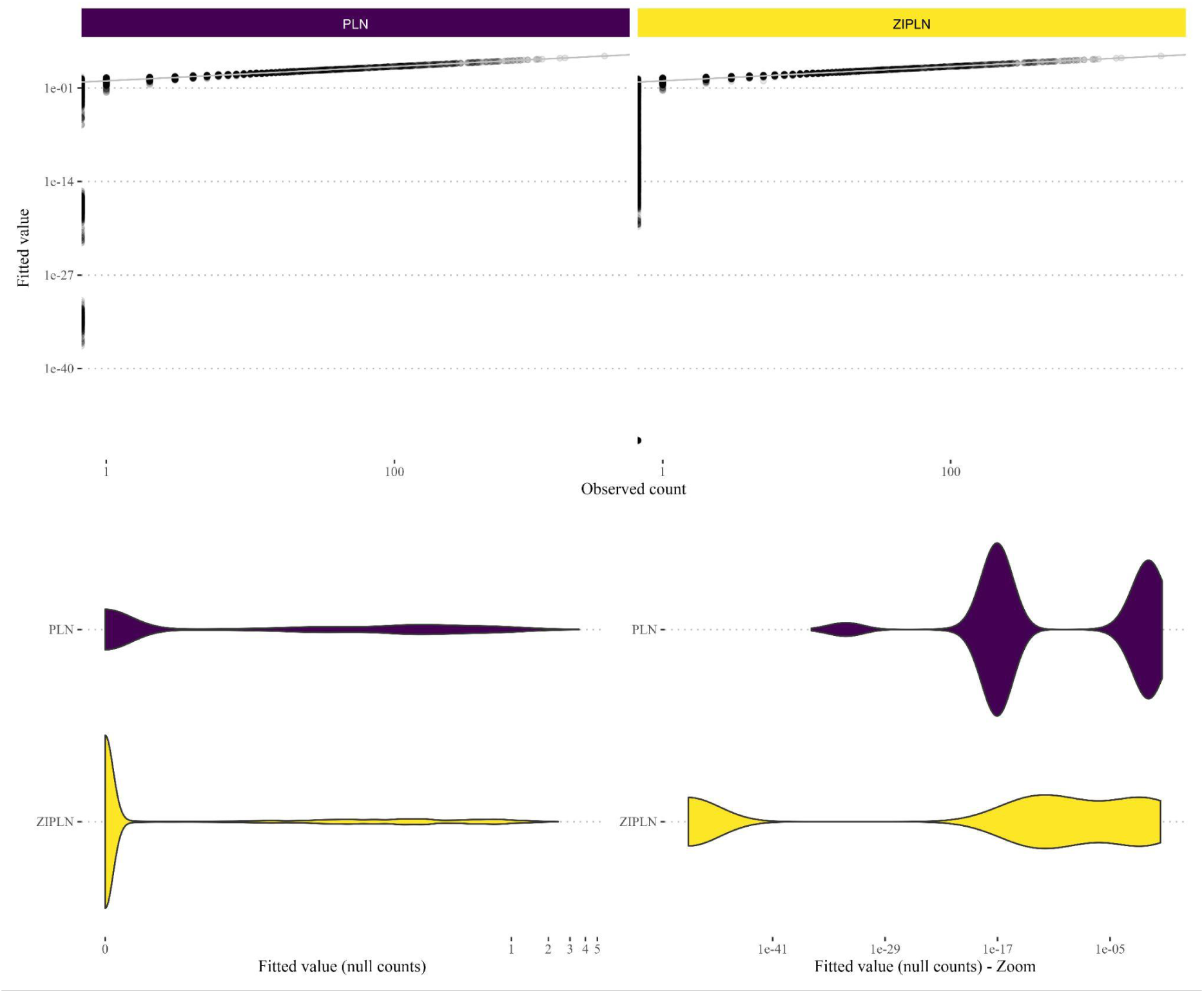
Model fit diagnostics for the PLN and ZIPLN models with the formula PLN-network(Y ∼ 1 + PC1 + PC2 + stream + disturbance) and ZIPLN-network(Y ∼ 1 + PC1 + PC2 + disturbance | 1 + stream). Fitted versus observed counts (left panel), fitted values for null counts (middle panel), fitted values for null counts with a zoom on the lowest fitted values (right panel).

